# Small molecule inhibitor of PPARγ acetylation promotes insulin sensitization and browning of white adipose tissue with improved safety

**DOI:** 10.64898/2025.12.15.694265

**Authors:** Dan Wu, Zhen Gong, Ying He, Sandeep Kumar, Venkateswararao Eeda, Qianfen Wan, Oana Herlea-Pana, Hui-Ying Lim, Wayne A. Hendrickson, Li Qiang, Weidong Wang

## Abstract

The nuclear receptor PPARγ is a primary therapeutic target for insulin resistance and type 2 diabetes (T2D); however, its thiazolidinedione (TZD) class of PPARγ agonists have substantial safety concerns in clinical utilization. Genetic inhibition of PPARγ acetylation at K268 and K293 dissociates the major adverse effects of TZDs from insulin sensitization and other metabolic improvements. We therefore posit that chemical inhibition of PPARγ acetylation would elicit insulin sensitization with improved safety. Here we describe the identification of a synthetic thiopyrimidine derivative (TPMD) that acts as a small molecule inhibitor of PPARγ acetylation. TPMD improves insulin sensitivity, promotes brown remodeling of white adipose tissue (WAT), increases energy expenditure, and decreases adiposity in dietary and genetic mouse models of T2D. Importantly, TPMD is deprived of the major side effects of TZDs, including weight gain, fluid retention, cardiac hypertrophy, bone marrow adiposity, and bone loss. X-ray crystallography at 2.0 Å resolution reveals a unique binding mode of TPMD molecules to the PPARγ ligand-binding domain and provides structural and molecular basis for the acetylation inhibitory activity of TPMD together with site-directed mutagenesis studies. These findings identify TPMD as a first-in-class compound that specifically targets PPARγ acetylation and possesses the potential of developing into a safe insulin sensitizer to treat and prevent T2D and obesity.

## Introduction

Insulin resistance is a key driving force for Type 2 diabetes (T2D) and other metabolic disorders. Moreover, although insulin resistance is considered as the primary comorbidity of obesity, it could develop much earlier than the onset of obesity. Restoring and improving insulin sensitivity is arguably the first strategy to prevent and treat T2D and its comorbidities. However, in contrast to the numerous medications to control hyperglycemia, there is only one class of drugs that directly targets insulin resistance - thiazolidinediones (TZDs). TZDs are the synthetic full agonists of PPARγ (Lazar, 2018), a nuclear receptor that is vital for insulin sensitivity, glucose and lipid metabolism, and adipocyte biology (Lehrke and Lazar, 2005; Spiegelman, 1998; Tontonoz et al., 1995; Tontonoz and Spiegelman, 2008). Unfortunately, the clinical utilizations of TZDs are curtailed by adverse side effects represented by weight gain, adiposity, edema, heart failure, and bone fractures. As such, there is an urgent unmet medical need of insulin sensitizer with improved safety.

PPARγ undergoes various post-translational modifications (PTMs) on multiple residues, including acetylation, phosphorylation, O-glycosylation, SUMOylation, and ubiquitination, for the regulation of protein abundance and function. Among the PTMs, acetylation at K268/K293 of PPARγ is heightened in adipose tissue under obesity and aging and shows a diurnal rhythm orchestrated with metabolic oscillation (He et al., 2023a; Qiang et al., 2012). Constitutive acetylation-mimetic of PPARγ K293 (K293Q) reduces adipose plasticity (He et al., 2023b) and worsens inflammation and adipose degeneration (Aaron et al., 2022) when over-expressed in adipocytes and in macrophages respectively. On the other hand, mice carrying the deacetylation mimetic of PPARγ (K268R/K293R, 2KR) retain the insulin-sensitizing effects of TZD but not the adverse side effects including bone loss, fluid retention, and heart hypertrophy when treated with the TZD drug rosiglitazone (Rosi) and are protected against diet-induced or aging-associated obesity (Kraakman et al., 2018). Collectively, the observations support a model that PPARγ deacetylation decouples the insulin-sensitization of TZD drugs from their side effects and provide a potential strategy to tackle insulin sensitivity with improved safety through targeting PPARγ acetylation. However, it is unclear whether PPARγ acetylation is targetable and sufficient on its own to confer the metabolic benefits while sparing the undesirable side effects.

In the present study, we sought to discover small molecules that modulate PPARγ acetylation. A synthetic thiopyrimidine derivative, 4-cyclohexyl-2-((2-nitro-4-(trifluoromethyl)phenyl)thio)-6-oxo-1,6-dihydropyrimidine-5-carbonitrile (termed as TPMD), was found to act as a small molecule modulator of PPARγ deacetylation. TPMD improves metabolic dysfunction related to diabetes and obesity; however, it is deprived of TZD’s major side effects. Moreover, TPMD promotes brown remodeling of white adipose tissue (WAT) and increases energy expenditure to protect against diet-induced obesity. X-ray crystallography resolves a unique binding pattern of TPMD to the PPARγ ligand binding pocket at a resolution of 2.0 Å, together with side-directed mutagenesis assays, providing structural and molecular basis for the deacetylation activity of TPMD. In sum, we identify TPMD as a first-in-class compound that promotes PPARγ deacetylation and possesses a potential of developing into a safe insulin sensitizer to treat and prevent T2D.

## Results

### Identification of TPMD as a novel synthetic ligand of PPARγ but with weak agonist activity

In our previous high throughput screening, we obtained a number of small molecules that prevent β cell dysfunction and death against endoplasmic reticulum (ER) stress (Duan et al., 2016a; Duan et al., 2016b; Duan et al., 2017; Tran et al., 2014). One hit compound, a synthetic derivative of thiopyrimidine (TPMD, Fig. 1A), protected the pancreatic β cell line INS-1 cells from ER stressor tunicamycin (Tm)-induced cell death (Fig. S1A-B). TPMD suppressed the cleavage (activation) of Tm-induced proapoptotic markers (Caspase 3 and poly ADP-ribose polymerase (PARP) (Fig. S1C) as well as Tm-induced ER stress markers (Fig. S1D-G) in INS-1 cells. Since PPARγ was reported to protect β cells against ER stress (Evans-Molina et al., 2009; Gupta et al., 2010), we investigated whether this compound or any other hit compound may act through modulating PPARγ activity. Among approximate 30 hits tested, TPMD was the only one that binds to PPARγ, with half-maximum inhibitory concentration (IC_50_) at ∼5 µM (Fig. 1B). TPMD is structurally distinct from all known PPARγ ligands, and its biological function has not been explored before. TPMD showed a dose-dependent activity on activating PPARγ, but much milder than the full agonist Rosi, in the classic 3xPPRE luciferase reporter assay (Fig. 1C). Similarly, TPMD also weakly activated PPARγ transcriptional activity on the endogenous *Adipsin* promoter-driven luciferase reporter (Fig. 1D).

**Figure 1:**
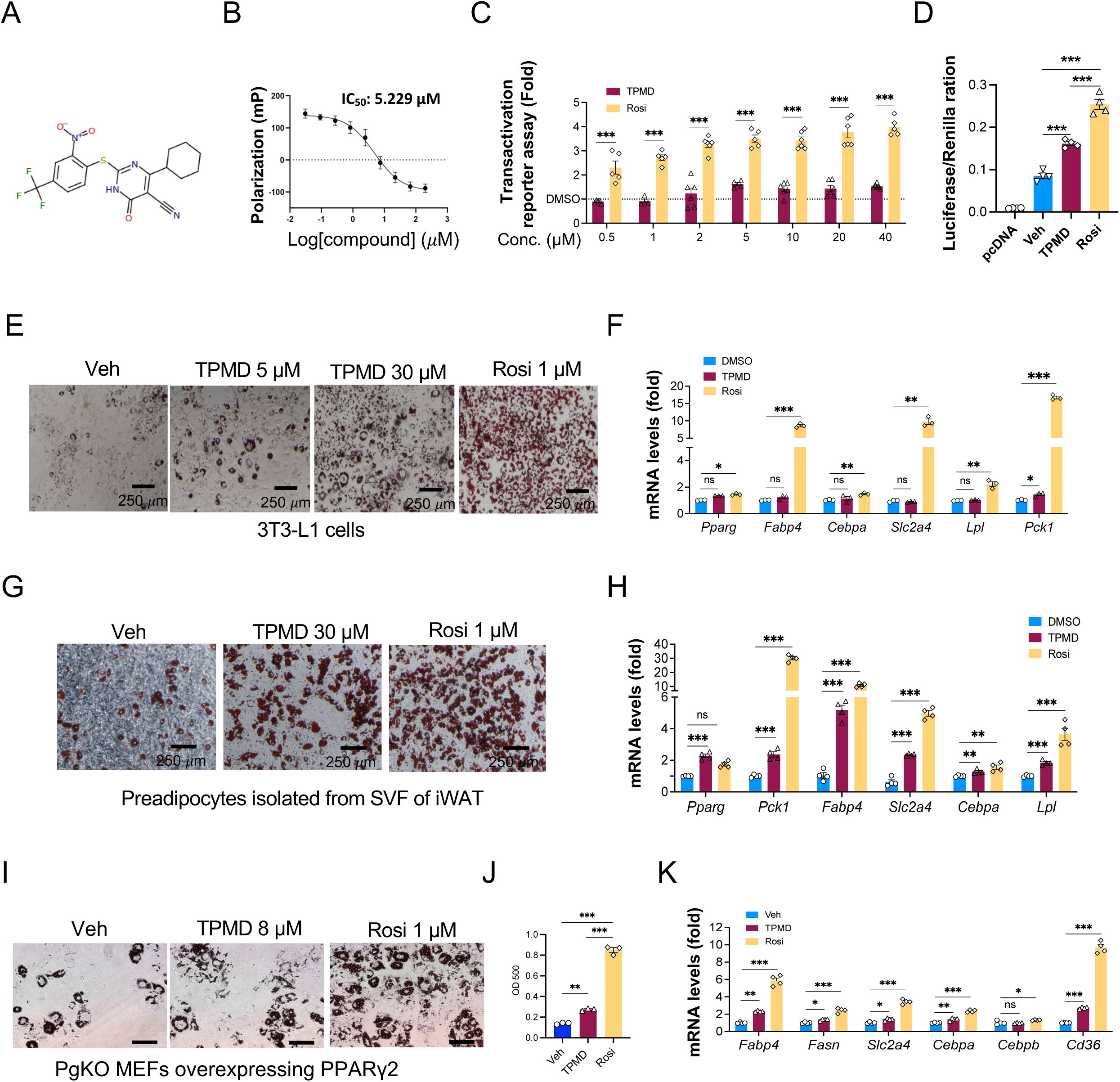
Identification of TPMD as a novel PPARγ partial agonist. A. Chemical structure of TPMD. B. *In vitro* competition binding assay between TPMD and purified PPARγ ligand-binding domain. Florescence polarization value (mP) was measured at 485 nm (excitation) and at 535 nm (emission). C. Transactivation assay for 3xPPRE luciferase reporter in HEK293 cells co-treated with indicated compounds for 24 hours. Fold changes of TPMD or Rosi over DMSO were presented. D. Transactivation assay for adipsin promoter luciferase reporter in HEK293 cells co-treated with indicated compounds for 24 hours. E-F. 3T3-L1 adipocyte differentiation under the indicated treatment of TPMD or Rosi for last 4 days; Oil Red O staining for lipid accumulation (E) and qRT-PCR analysis of relative mRNA expression levels of adipogenic genes (F). G-H. Primary mouse adipocyte differentiation with SVF isolated from iWAT under the indicated treatment of TPMD or Rosi for last 4 days; Oil Red O staining for lipid accumulation (G) and qRT-PCR analysis of relative mRNA expression levels of adipogenic genes (H). I-K. Oil Red O staining (I) of PPARγ KO MEFs that were engineered to constitutively express PPARγ differentiated under the indicated treatment of TPMD or Rosi for last 4 days, OD 500 reading of the staining (J), and qRT-PCR analysis of relative mRNA expression levels of pan adipocyte marker genes (K). Data are the mean± SEM and are representative of 3 independent experiments. * P < 0.05, ** P <0.01, and *** P <0.001. ns for no significance.

PPARγ is the master regulator of adipogenesis. Next, we adopted adipocyte differentiation as a functional test of TPMD activity. TPMD dose-dependently promoted lipid accumulation during 3T3-L1 cell differentiation, but significantly less than Rosi even at a 30-fold higher dose (Fig. 1E, S2A). At the molecular level, TPMD exhibited much weaker activity in inducing adipogenic genes than Rosi, many of which are PPARγ downstream targets, including *Fabp4, Pparg, Cebpa*, *Slc2a4 (*also known as *Glut4), Lpl,* and *Pck1* (Fig. 1F). This weak adipogenic effects of TPMD was recapitulated in primary mouse adipocyte differentiation in terms of lipid accumulation and adipogenic gene expression (Fig. 1G-H, S2B-C). Further, to exclude the possibility that TPMD acts through the induction of PPARγ transcription to facilitate adipogenesis, we employed the PPARγ knockout mouse embryonic fibroblasts (MEFs) reconstituted with constitutively expressed wildtype PPARγ2 (PgKO-PPARγ2) (Li et al., 2016). Again, TPMD also showed a weaker adipogenic activity in these cells (Fig. 1I-K), supporting the notion that TPMD acts on PPARγ directly rather than through its induction. Therefore, we have identified TPMD as a new scaffold that binds to PPARγ but acts as a weak agonist.

### TPMD improves insulin sensitivity and other metabolic dysfunction in diet-induced obesity

Next, we evaluated the effects of TPMD on metabolic dysfunctions in mouse diet-induced obesity (DIO) model (Fig. 2A). Four week’s treatment of TPMD significantly ameliorated insulin resistance in DIO mice (Fig. 2B), accompanied by marked decrease in fasting plasma insulin levels (Fig. 2C). Glucose tolerance was also improved (Fig. 2D). Fittingly, TPMD increased the expression of a set of PPARγ downstream target genes that were reported to be associated with insulin sensitivity (Choi et al., 2010), including *adiponectin* (*adipoq*)*, adipsin, selenbp1, Rarres2*, and *Car3*, in the eWAT (Fig. S3A) and iWAT (Fig. S3B). In contrast, the adipogenic house-keeping genes were either even slightly down-regulated in eWAT (Fig. S3C) or essentially unaltered in iWAT (Fig. S3D) in TPMD-treated DIO mice. Moreover, TPMD improved DIO-associated dyslipidemia, decreasing triglyceride (TG), free fatty acids (FFA), and total cholesterol levels in circulation (Fig. 2E-G). Hepatic steatosis was also alleviated by TPMD treatment (Fig. 2H), with accompanying downregulation of key lipogenic and gluconeogenic genes in liver tissues (Fig. S3E, F). Therefore, despite as a weak ligand of PPARγ, TPMD displays a potent insulin-sensitizing effect and substantially corrects the metabolic dysfunctions in DIO mice.

**Figure 2:**
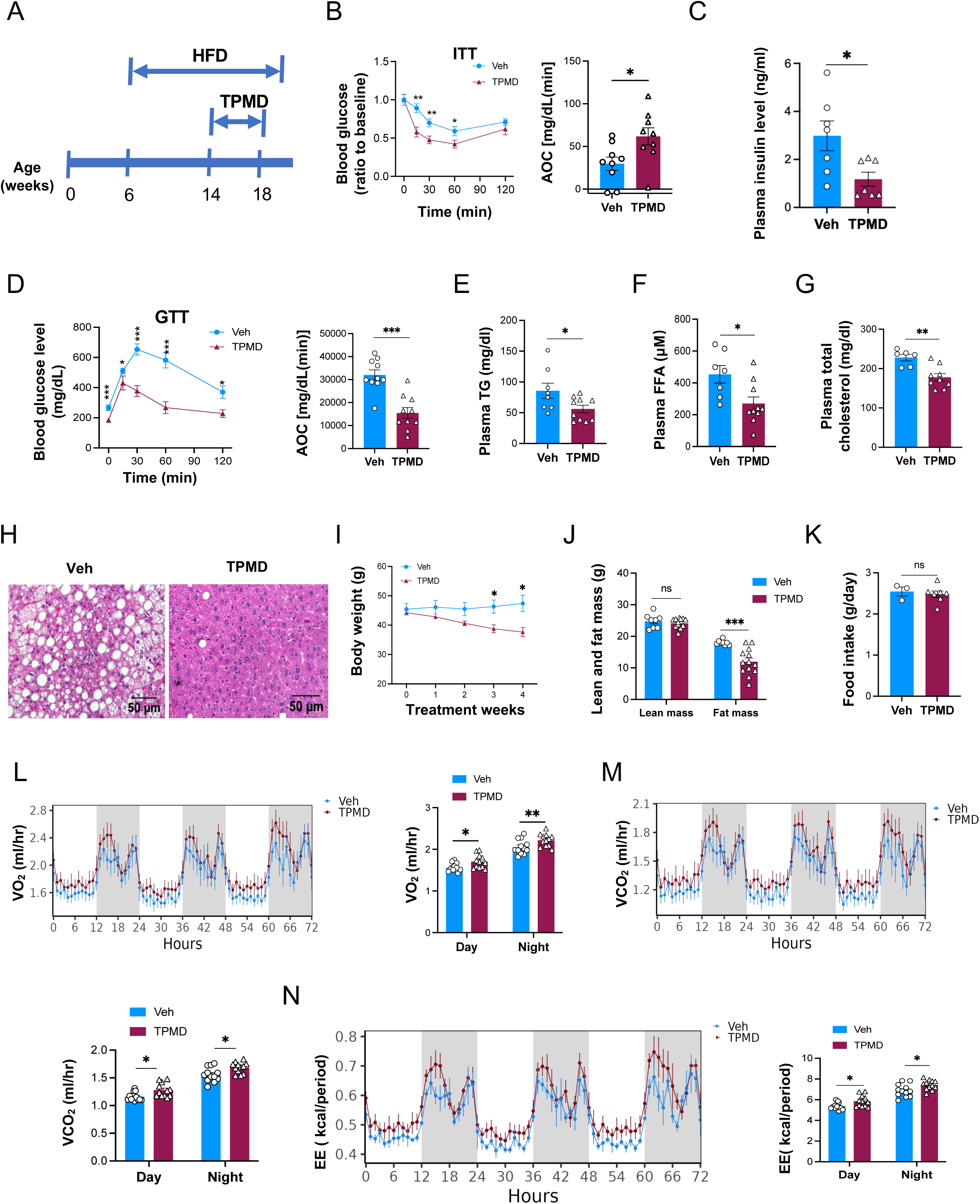
TPMD improves insulin sensitivity and energy expenditure and protects against diet-induced obesity and comorbidity. A. Timing of obesity induction and i.p. drug administration. B. Insulin tolerance test performed for mice treated with TPMD (n=10) or vehicle (n=10) for 4.5 weeks. Blood glucose levels normalized to basal level at indicated time points after intraperitoneal injection of insulin (1.0 IU/kg body weight) following 6-h fasting and the AOC (area of the curve). C. Plasma insulin levels following 6-h fasting. D. Glucose tolerance test performed for mice treated with TPMD or vehicle for 4 weeks. Blood glucose levels measured at indicated time points after intraperitoneal injection of glucose (1.5 g/kg body weight) following 14-h fasting and the AOC. E. Plasma total cholesterol level. F. Plasma free fatty acid level. G. Plasma triglyceride level. H. H&E staining of liver sections from DIO mice treated with TPMD or vehicle for 4.5 weeks. I. Body weight of DIO mice treated with TPMD or vehicle. J. Fat and lean mass by EchoMRI. K. Daily food intake, measured during the 4^th^ week of treatment. L-M’. Metabolic cage analysis of DIO mice treated with TPMD or vehicle after 4-week treatment, Measurement of oxygen (L) and carbon dioxide (M) consumption levels. N. Energy expenditure levels. Data are the mean± SEM and are representative of 3 independent experiments. * P < 0.05, ** P <0.01, and *** P <0.001.

Besides imposing the potent insulin-sensitizing activity, TPMD also reduced body weight in DIO mice, resulting in 16% less than the vehicle-treated control mice after 4 weeks’ treatment (Fig. 2I). This body weight reduction was exclusively contributed by decrease in fat mass without change in lean mass (Fig. 2J). This anti-obesity effect was not caused by reduced food intake (Fig. 2K). Instead, TPMD treatment increased oxygen consumption, CO_2_ production, and energy expenditure in DIO mice during both the light and dark phases by indirect calorimetry analysis (Fig. 2L-N, S3G-I) without affecting locomotion activity and respiratory exchange ratio (RER) (Fig. S3J, K-L).

### TPMD improves adipose tissue remodeling in obesity

Underlying the reduced fat mass is the alleviation of adipocyte hypertrophy in both eWAT (Fig. 3A, S4A) and iWAT (Fig. 3B, S4B) in TPMD-treated DIO mice. A pervasive feature of obesity is adipose tissue inflammation indicated by macrophage infiltration (Hotamisligil et al., 1995), which correlates with adipocyte hypertrophy (Weisberg et al., 2003). The crown-like structures formed by macrophages were remarkably reduced in the eWAT of TPMD-treated DIO mice compared to vehicle-treated mice (Fig. 3A, C). Consistently, inflammatory genes *Tnfa*, *Il6*, *Il1b*, and *Ccl2* were also repressed (Fig. 3D), accompanied by attenuated expression of macrophage markers *Adgre1* (also known as *F4/80*) and *Cd68* (Fig. 3E), and a diminishment in the accumulation of F4/80-positive cells (Fig. S4C, D), with no repressive effect on anti-inflammatory genes *Il10* and *Mgl1* (Fig. S4E), in the eWAT from TPMD-treated DIO mice. Therefore, TPMD mitigates the pathological remodeling of adipose tissue in obesity.

**Figure 3:**
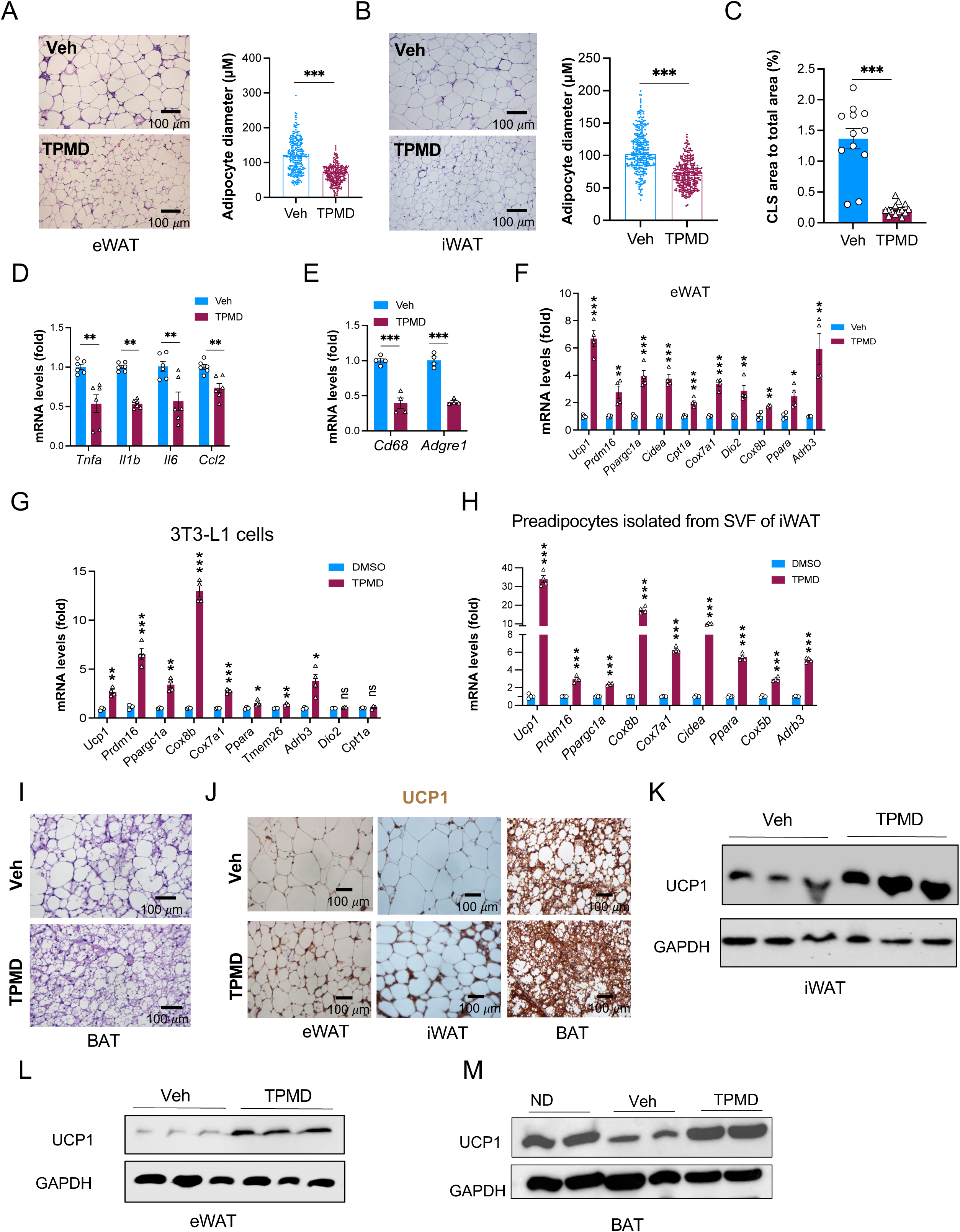
TPMD promotes adipose tissue remodeling. A. H&E staining of eWAT sections from DIO mice treated with TPMD or vehicle; representative images of eWAT and average diameter of adipocytes (μM)/field. B. H&E staining of iWAT sections form DIO mice treated with TPMD or vehicle; representative images of of iWAT and average diameter (μM)/field. C. CLS area measurement in eWAT from DIO mice treated with TPMD or vehicle for 4 weeks. D-E. qRT-PCR analysis of mRNA levels of genes for M1 proinflammatory markers (D) and macrophage markers (E). F. qRT-PCR analysis for mRNA levels of genes known for browning/thermogenesis in eWAT from DIO mice treated with TPMD or vehicle. G. qRT-PCR analysis for mRNA levels of browning/thermogenic marker genes in adipocytes differentiated from 3T3-L1 cells under the indicated treatment of TPMD or DMSO control for last 4 days. H. qRT-PCR analysis for mRNA levels of browning/thermogenic marker genes in adipocytes (differentiated from wild-type iWAT) under the indicated treatment of TPMD or DMSO for last 4 days. I. H&E staining of BAT from DIO mice treated with TPMD or vehicle. J. Immunohistochemical staining of UCP1 in eWAT, iWAT, and BAT sections from DIO mice treated with TPMD or vehicle. K-M. Immunoblotting of protein levels of UCP1 in eWAT, iWAT, and BAT from DIO mice treated with TPMD or vehicle. GAPDH as loading control. Data are the mean± SEM. * P < 0.05, ** P <0.01, and *** P <0.001.

In line with the increased energy expenditure, TPMD treatment upregulated browning genes, including *Ucp1*, *Prdm16*, *Pgc1a, Cidea*, *Cpt1a, Cox8b, Cox7a1*, and *Trem26*, in both eWAT and iWAT (Fig. 3F, S4F). This increase of browning genes was also observed *in vitro* in adipocytes differentiated from 3T3-L1 cell line (Fig. 3G) and in primary adipocytes (Fig. 3H, S4G), suggesting a cell-autonomous browning effect. Moreover, BAT undergoes a whitening process featured by lipid accumulation in obesity. Histological analyses of BAT revealed a strong inhibition of lipid accumulation in DIO mice by TPMD (Fig. 3I). Further, the brown adipocyte marker UCP1 was increased by TPMD treatment, as shown by immunohistochemical staining (Fig. 3J) and western blotting (Fig. 3K-M) in iWAT, eWAT, and BAT. In addition, the frequency in the occurrence of multilocular adipocytes (a hallmark feature of beige adipocyte) in the WAT of DIO mice was elevated in TPMD-treated mice (Fig. S4H). TPMD therefore improves adipose tissue remodeling in DIO by promoting the brown remodeling of WAT and reversing the whitening of BAT.

### TPMD displays the similar metabolic benefits in *ob/ob* mice

We then interrogated whether the metabolic effects of TPMD was restricted to DIO model. To this end, we treated mice of leptin deficiency (*ob/ob*), a genetic obesity model, with TPMD. Similarly, TPMD improved glucose tolerance and insulin sensitivity (Fig. S5A, 4A-B) and decreased fasting plasma insulin levels (Fig. 4C). TPMD-treated *ob/ob* mice for 4 wks exhibited approximately 30% weight gain as their vehicle-treated counterparts (Fig. 4D) without changes in food intake (Fig. 4E). Likewise, the alleviation of adipocyte hypertrophy by TPMD observed in DIO was recapitulated in *ob/ob* mice (Fig. 4F). In addition, we observed that TPMD treatment elevated the expression of insulin sensitivity-related genes in both eWAT and iWAT (Fig. 4G, S5B), whereas it either reduced or had little effect on the expression of adipogenic house-keeping genes (Fig. 4H, S5C). TPMD treatment also suppressed the accumulation of F4/80^+^ macrophages in the eWAT (Fig. 4I, J) and iWAT (Fig. S5D, E) of *ob/ob* mice. Furthermore, TPMD treatment ameliorated dyslipidemia in *ob/ob* mice (Fig.4K-M). Finally, liver steatosis was also markedly ameliorated in TPMD-treated *ob/ob* mice (Fig. 4N, O). Collectively, these data indicate that TPMD exerts efficient insulin sensitization and metabolic improvements in both obesity models. In contrast, when administered to chow-fed mice, TPMD did not exhibit noticeable effects except a moderate improvement of iPGTT in TPMD-treated animals (Fig. S6A-J).

**Figure 4:**
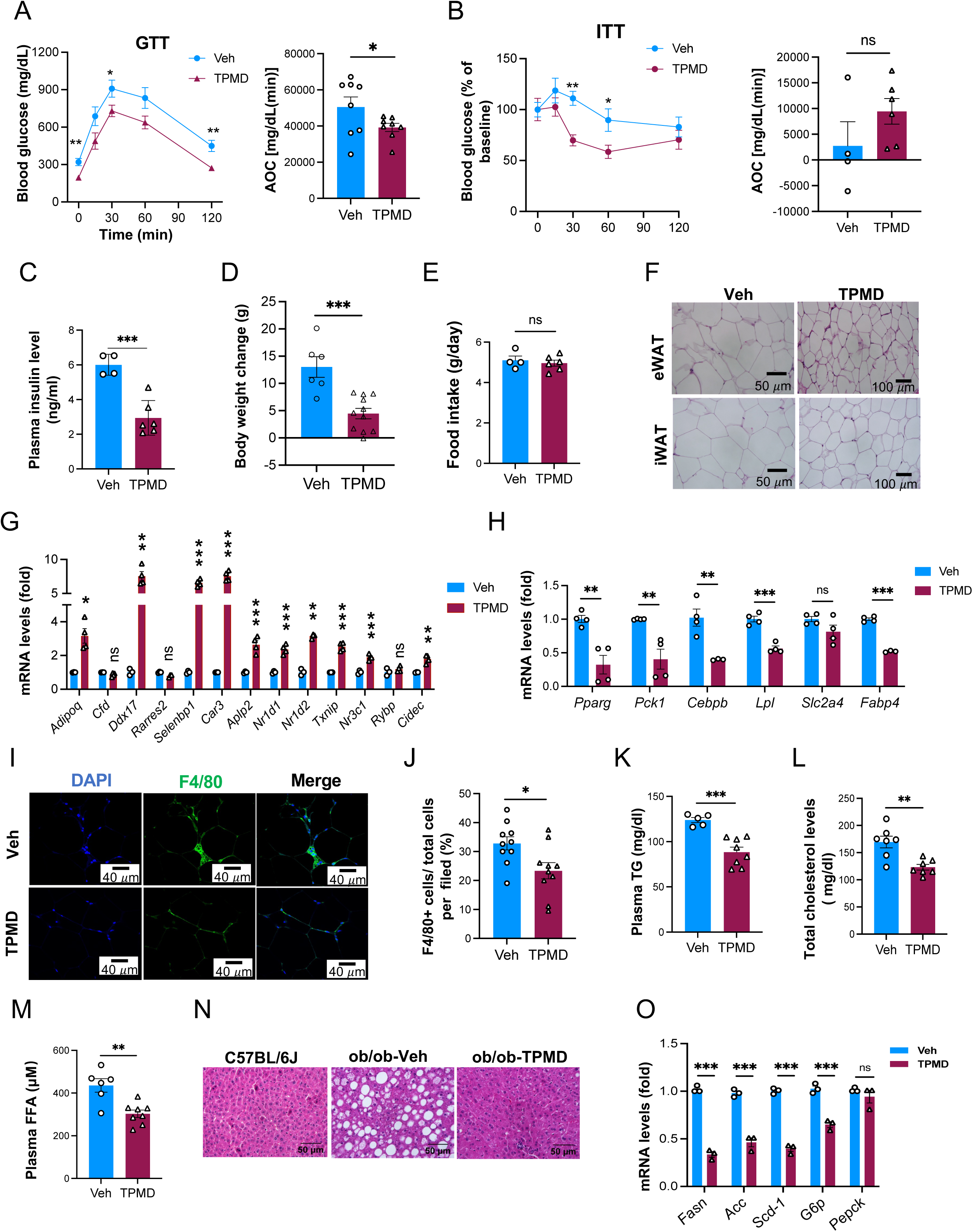
TPMD improves insulin sensitivity and comorbidity in Leptin-deficient obese mice. A. Glucose tolerance test from ob/ob mice treated with TPMD or vehicle for 4 weeks. Blood glucose levels measured at indicated time points after intraperitoneal injection of glucose (1.5 g/kg body weight) following 14-h fasting and the AOC. B. Insulin tolerance test performed for mice treated with TPMD or vehicle for 4.5 weeks. Blood glucose levels normalized to basal level at indicated time points after intraperitoneal injection of insulin (1.0 IU/kg body weight) following 6-h fasting and the AOC. C. Plasma insulin levels fasted overnight of mice treated with TPMD or vehicle for 4 weeks. D. Changes in body weight of ob/ob mice treated with TPMD or vehicle over the period of 4-wk treatment. E. Daily food intake, measured for three consecutive days during the 4^th^ week of treatment. F. H&E staining of eWAT and iWAT sections from ob/ob mice treated with TPMD or vehicle. G-H. qRT-PCR analysis for mRNA levels of genes known for insulin sensitivity (G) and for adipocyte housing-keeping (H) in eWAT from ob/ob mice treated with TPMD or vehicle. I-J. Immunofluorescent staining of F4/80 in eWAT sections from ob/ob mice treated with TPMD or vehicle for 4 weeks; representative images (I) of F4/80 staining and quantification of percentage of F4/80 positive cells (J). K. Plasma triglyceride level. L. Plasma total cholesterol level. M. Plasma free fatty acid level. N. H&E staining of liver sections from ob/ob mice treated with TPMD or vehicle for 4.5 weeks. O. qRT-PCR analysis of mRNA levels of lipogenic and gluconeogenic markers in liver tissues from ob/ob mice treated with TPMD or vehicle. Data are the mean± SEM. * P < 0.05, ** P <0.01, and *** P <0.001.

### TPMD is devoid of adverse side effects associated with TZD use

The major clinical obstacle of TZDs is their adverse side effects, primarily weight gain, fluid retention, cardiac risk, and bone loss (Nanjan et al., 2018). In contrast to TZDs, TPMD showed an anti-obesity effect by reducing adiposity as shown above (Figs. 2I, J, 3A, C). Rosi caused fluid retention (hemodilution) in DIO mice, as reflected by a decrease in the hematocrit level (PCV%), whereas TPMD showed little effect (Fig. 5A). In contrast to TZD treatment which is associated with cardiac hypertrophy (Khalaf and Taegtmeyer, 2012), 4-wk TPMD treatment did not increase heart weight (Fig. 5B, C), underpinned by the blunted induction of cardiac genes associated with heart failure or hypertrophy, including myosin heavy chain β (β*-Mhc,* also *Myh7*), natriuretic peptide b (*Nppb*), *Bnp*, and *Acta1* (Fig. 5D) (Choi et al., 2011). Moreover, Rosi upregulated genes associated with lipotoxicity in the heart, such as *Fabp4*, *Cd36*, *Acacb*, *Srebf1,* and *Bscl2*; however, no such upregulation was observed after TPMD treatment (Fig. 5E). Furthermore, while there was no noticeable difference in bone density between Rosi and TPMD treatments of 4 weeks (Fig. 5F), Rosi treatment downregulated the expression of bone formation genes in the tibia of DIO mice, in contrast to the blunted effect of TPMD (Fig. 5G). Bone marrow adiposity (BMA) is tightly correlated with bone loss (Ghali et al., 2016). Rosi promoted BMA expansion whereas TPMD displayed no effect on BMA expansion (Fig. 5H, I). Consistently, unlike Rosi, TPMD treatment did not induce adipogenic genes (*Pparg2*, *Cfd*, *aP2*, *Perilipin*, *Adipoq*) (Fig. 5J) or repress osteogenic genes (*Osteocalcin*, *Col1a1*, *Alp*, *Spp1*) (Fig. 5K) in the bone marrow of DIO mice. Together, these results indicate that TPMD is spared of the major side effects associated with TZDs.

**Figure 5:**
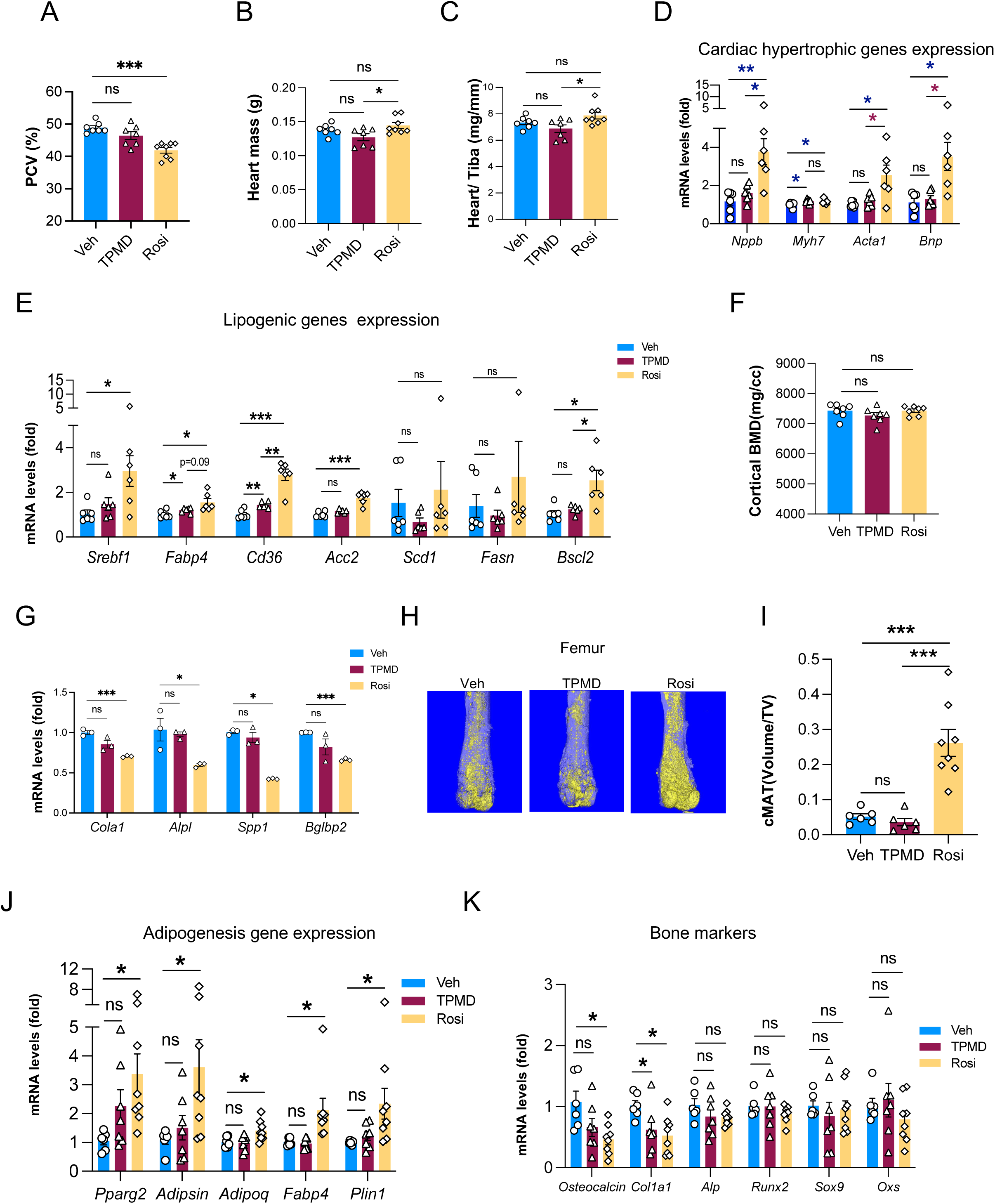
TPMD is devoid of the adverse side effects associated with TZD use. A. Measurement of PCV in DIO mice treated with TPMD, Rosi, and vehicle. B. Heart weight of DIO mice treated with TPMD, Rosi, or vehicle. C. Ratio of heart weight relative to the length of tibia bone (mg/mm). D-E. qRT-PCR analysis of mRNA levels of heart failure/hypertrophy associated genes (D) and lipogenic genes (E) in hearts from DIO mice treated with TPMD or vehicle. F. Femoral bone mineral density in the cortical regions by μCT scanning. G. qRT-PCR analysis of mRNA levels of bone density/formation-associated genes in bone tissue from DIO mice treated with TPMD, Rosi or vehicle. H-I. Osmium tetroxide staining of femoral marrow lipid droplets (areas of golden color) (H) and quantification (I) by μCT scanning. J-K. qRT-PCR analysis of mRNA levels of adipogenic genes (J) and bone marker genes (K) in femoral bone marrow from DIO mice treated with TPMD or vehicle. Data are the mean± SEM. * P < 0.05, ** P <0.01, and *** P <0.001.

### Crystallography reveals a unique binding mode of TPMD to PPAR**γ**

To interrogate the molecular basis of the functional profile of TPMD on PPARγ, we determined the crystal structure of PPARγ ligand binding domain (LBD) in complex with TPMD by X-ray crystallography at a resolution of 2.0 Å (PDB code: 9ZL6). The detailed statistics of diffraction data collection and structure refinement are summarized in Table S1. Unexpectedly, the crystal structure reveals that a single PPARγ LBD accommodates three TPMD molecules, each adopting a distinct binding mode (Fig. 6A). Excellent electron density coverage was obtained for all three bound TPMD molecules (Fig. S7A and B), from which their conformations could be unambiguously determined. The PPARγ ligand binding pocket is a large hydrophobic T-shaped cavity containing three branches (Kroker and Bruning, 2015a; Nolte et al., 1998). Two TPMD molecules are located within the ligand-binding pocket, with one TPMD molecule (TMPD-1) located in the left branch and the other TPMD molecule (TMPD-2) positioned in the right branch. The third TPMD molecule (TPMD-3) resides in a cleft formed by helices H3, H4, H5 and H12 (Fig. 6A).

**Figure 6:**
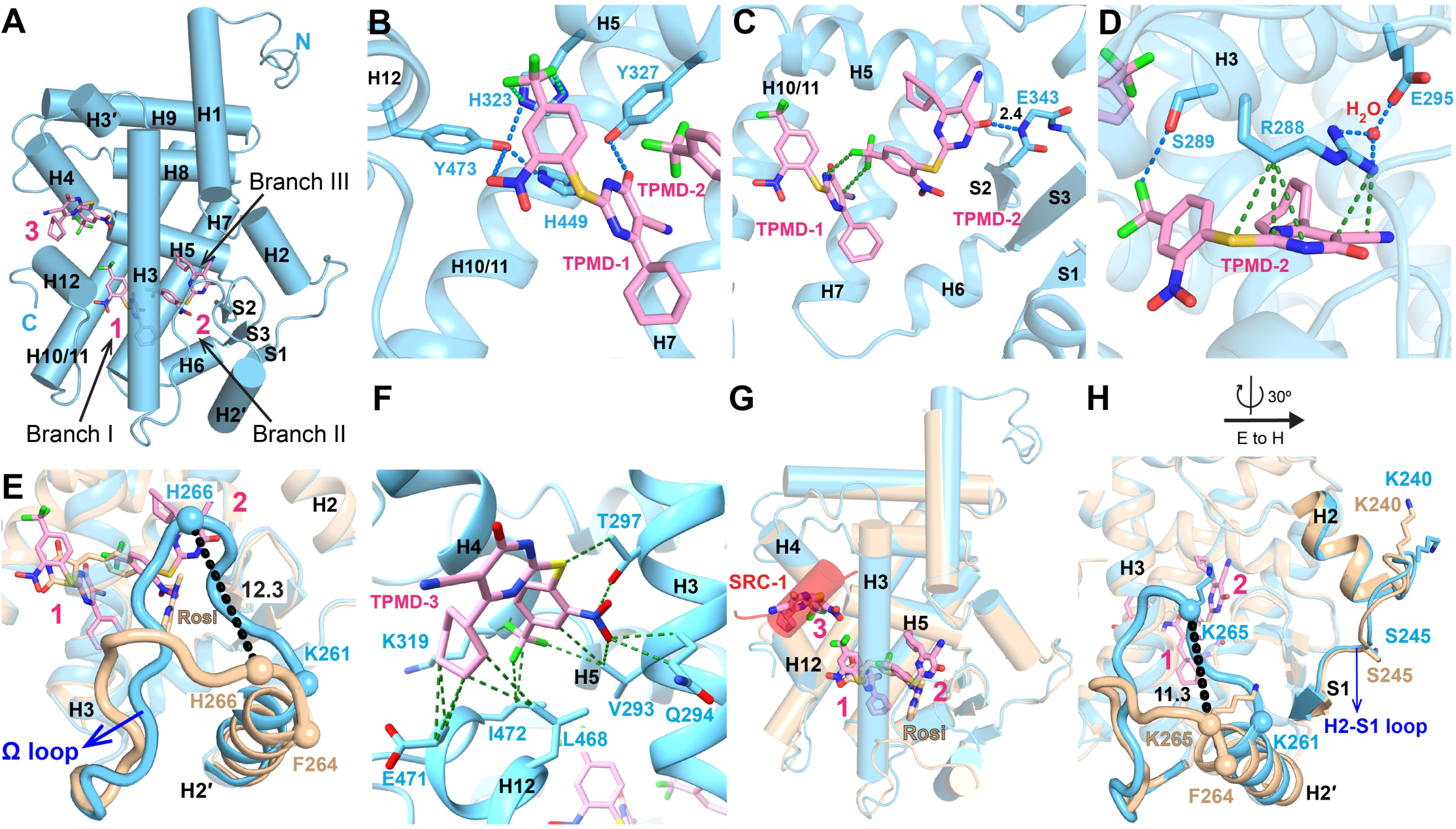
Structure of the PPARγ LBD in complex with TPMD. **(A)** An overall structure of the PPARγ LBD, amino acids 223-505, in complex with TPMD. The LBD is shown in light blue cartoons while TPMDs are shown in stick representation with carbon, nitrogen, oxygen and fluorine atoms depicted in pink, blue, red and green respectively. (**B**) Interactions between TPMD-1 and PPARγ LBD. The protein residues in TPMD-bound PPARγ LBD are shown in stick representation with carbon atoms depicted in light blue. (**C**) Interactions between TPMD-2 and PPARγ LBD as well as interactions between TPMD-1 and TPMD-2. (**D)** Interactions between TPMD-2 and PPARγ LBD. (**E**) A comparison of the Ω loop in the TPMD-bound (blue) and Rosi-bound (beige) PPARγ LBD. The Cα atoms of F292 are shown as spheres in each to highlight the Ω loop displacement. The Cα atoms of K289 and H294, also shown as spheres, mark the C-termini of each H2′ helix to highlight unwinding for the TPMD complex. (**F)** Interactions between TPMD-3 and PPARγ LBD. (**G**) Superimposition of the TPMD-bound (light blue) and Rosi-bound (beige) PPARγ LBD. **(H)** Conformational changes of K268 and K293 in the TPMD-bound PPARγ LBD (shown in light blue ribbons and sticks) compared with Rosi-bound LBD (shown in beige ribbons and sticks). Blue, green and black dashes represent H bonds, favored vdW contacts and distance between atoms. The Ω loop is highlighted using thicker loop. Helix 3 was hidden for better display of TPMDs in Figure B and C.

TPMD-1 resides near the left branch (branch I) composed of a mix of hydrophobic residues and polar residues from α helices H3, H5, H10/11 and H12. It adopts a linear conformation extending below the N terminus of helix 5 to the bottom half of helix 3 by making several specific interactions with residues in this branch (Fig. 6A and B). The nitro group of TPMD-1 forms a hydrogen bond with the side chain hydroxy group of Y501 in helix 12 (Fig. 6B) while the carbonyl group in the pyrimidine ring of TPMD-1 makes another hydrogen bond with the side chain hydroxy group of Y355 in helix 5 (Fig. 6B). Additionally, the hydroxyl group of Y501 in helix 12 also hydrogen bonds with H351 in helix 5 and with H477 in helix 10/11 (Fig. 6B). Branch I of the pocket is the canonical binding site for the acidic head group of TZD agonists (Nolte *et al*., 1998). Rosi forms hydrogen bonds directly with H351 and H477 and a secondary hydrogen bond with Y501 (Nolte *et al*., 1998), allowing for stabilization of the AF-2 surface (formed by helices H3, H4, H5 and H12 of PPARγ LBD) which is critical for co-activator recruitment and for full agonism of PPARγ. However, the conformation of these residues is altered in the TPMD-bound complex (Fig. S7C). Notably, F391, which shows high B factors and is presumably mobile in the apo structure and several ligand-bound structures (Itoh et al., 2008), is repositioned to avoid clashes with TPMD-1 (Fig. S7D).

TPMD-2 in the right branch (branch III) adopts a U-shaped conformation and is positioned between the lower half of H3 and the β sheets (Fig. 6A and C). This region is part of an alternative binding site that is likely critical for PPARγ-dependent insulin sensitization (Hughes et al., 2014). TPMD-2 makes several specific interactions with amino acids in H3 and the β turn between β2 and β3 strands. The carbonyl group in the pyrimidine ring of TPMD-2 forms a strong hydrogen bond (2.4 Å) with the backbone amide of E371 at the β turn between β2 and β3 strands (Fig. 6C). The CF_3_ group of TPMD-2 forms a weak hydrogen bond (3.2 Å) with the hydroxy side chain of S317 in H3 (Fig. 6D). TPMD-2 makes substantial favored van der Walls (vdW) contacts with the side chain of R316 of H3, which is hydrogen-bonded with the side chain of E323 through a water molecule (Fig. 6D). The side chains of R316, S317, E323, and E371 have substantial conformational changes to accommodate interactions with TPMD-2 compared with Rosi (Nolte *et al*., 1998) (Fig. S7D, E). Interestingly, the C terminus of H2′ is unwound almost one α-helix turn (residues 290-292) to move toward TPMD-2. As a consequence, the cylinder axis of H2’ moves 12° inward into the binding pocket. The Cα of H294 in the Ω loop moves 12.3 Å toward TPMD-2 relative to Rosi-bound PPARγ LBD (Nolte *et al*., 1998) (Fig. 6E). As so displaced, the Ω loop has made several favored vdW contacts with TPMD-2 and the β turn between β2 and β3 strands. For instance, F292 in the Ω loop are within favored vdW contacts to TPMD-2 (Fig. S7F). I290 and F292 in the Ω loop make favored vdW interactions with Q373 and S370 in the β turn between strands β2 and β3 respectively (Fig. S7F and G). Although the Ω loop of TPMD-bound PPARγ LBD could be reasonably traced by the electron density, the average B factor of the Ω loop is still higher than the overall B factor of the PPARγ LBD, indicating a certain degree of mobility of the Ω loop. Another key feature is that TPMD-2 extends deeply into branch III of the ligand binding pocket to interact with R316 and S317 in H3 and E371 in the β turn between β2 and β3 strands while in vicinity with the Ω loop simultaneously. This is very different from partial agonists SB1495 (Jang et al., 2019b) and Imatinib (Jang et al., 2019c), which stabilize the Ω loop by stretching to the opening of branch II of the ligand-binding pocket, or from partial agonist VPS-77 which suppresses PPARγ S273 phosphorylatio and interacts with the middle part of the alternative binding pocket (Jiang et al., 2020). In addition, TPMD-2 makes interactions with TPMD-1 through favored vdW interactions between the CF_3_ group of TPMD-2 and the pyrimidine ring of TPMD-1 (Fig. 6C).

TMPD-3 is located in a groove of the AF-2 surface through favored vdW interactions with V321, Q322 and T325 in H3, K347 in H4, and L496, E499 and I500 in H12 (Fig. 6A and 6F). TMPD-3 has occupied the position of the SRC-1 LXXLL motif in a Rosi-bound PPARγ LBD co-crystallized with SRC-1 LXXLL containing peptide (Fig. 6G), even though SRC-1 LXXLL was included in a 2:1 molar ratio to the PPARγ LBD during crystallization. We also co-crystalized the PPARγ LBD with the SRC-1 LXXLL motif-containing peptide in the absence of TPMD molecules and determined the structure at 2.2 Å resolution (PDB code: 9ZL5). The LXXLL motif then occupies the coactivator binding site on the AF-2 surface as in the Rosi-bound structure (Fig. S7H). These observations suggest that TPMD molecules compete with the co-activator binding site of the PPARγ LBD, which may result in weakened activation of the receptor. It is also worth mentioning that TMPD-3 resides near the N terminus of helix 1 (H1^s^) from a neighboring PPARγ LBD in the crystal lattice (Fig. S7I). To the best of our knowledge, it is the first time that the SRC-1 LXXLL motif is found to be displaced by a small molecule from a PPARγ LBD/SRC-1 LXXLL co-crystallization complex. Collectively, our structural data indicate that TPMD binds to PPARγ LBD in a mode distinct from that of full agonists (e.g., Rosi), partial agonists, or ligands that inhibit PPARγ S273 phosphorylation, which may underpin its unique *in vivo* functional profile.

### TPMD induces deacetylation of PPAR**γ**

The effects of TPMD on insulin sensitivity and safety profile largely phenocopy those seen in mice carrying the PPARγ deacetylation mimetics (K268R/K293R, 2KR) treated with TZD (Kraakman *et al*., 2018; Liu et al., 2020). We therefore tested whether TPMD impacts PPARγ deacetylation. Indeed, we found that TPMD facilitated the deacetylation of PPARγ2 (Fig. 7A) and at K268 and K293 (Fig. 7B) in HEK293 cells co-transfected with PPARγ, the acetyltransferase CREB-binding protein (CBP), and the deacetylase SirT1. As expected, TPMD also induced the *in vivo* PPARγ deacetylation in eWAT from TPMD-treated DIO mice (Fig. 7C). We further confirmed the deacetylation effect of TPMD on PPARγ using an *in vitro* assay where immunoprecipitation-purified acetylated PPARγ protein was incubated with recombinant Sirt1 protein in the presence of TPMD (Fig. 7D). Moreover, considering the positive correlation of K293 acetylation with phosphorylation on S273 (Qiang *et al*., 2012), we observed that TPMD inhibited S273 phosphorylation of PPARγ *in vitro* and *in vivo* (Fig. S8A, B). To further investigate the activity of TPMD on PPARγ acetylation, we utilized the P467L PPARγ mutant which is defective in binding to SirT1 and thus hyperacetylated (Qiang *et al*., 2012), to assess transcriptional activity. In contrast to Rosi, TPMD showed blunted activation of P467L mutant on 3xPPRE and *Adipsin* promoter-driven luciferase reporters (Fig. 7E, F). These results together suggest that TPMD elicits deacetylation of PPARγ.

**Figure 7:**
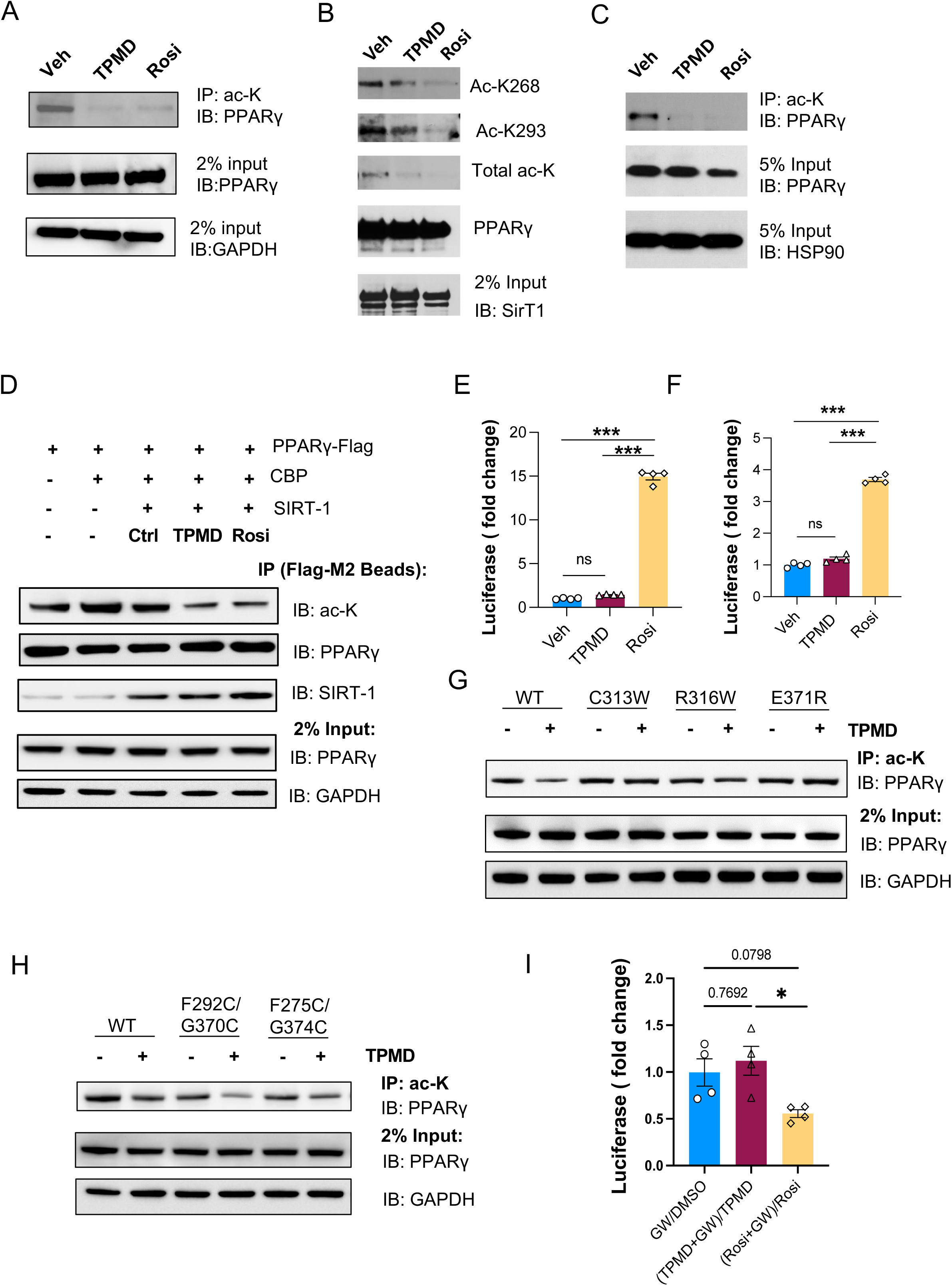
TPMD promotes PPARγ deacetylation. A. PPARγ acetylation in HEK293 cells co-transfected with PPARγ, acetyltransferase CBP, and deacetylase SirT1, in the presence of TPMD (30 μM) or Rosi (10 μM) as indicated. Total protein lysates were immunoprecipitated (IP) by acetylated lysine (ac-K) antibody conjugated agarose beads and IB for PPARγ. Rosi as a positive control. GAPDH was used as a loading control of total protein input. B. PPARγ acetylation in HEK293 cells co-transfected with Flag-tagged PPARγ, acetyltransferase CBP, and deacetylase SirT1, in the presence of TPMD (30 μM) or Rosi (10 μM) as indicated. PPARγ was immunoprecipitated by Flag M2 agarose beads and immunoblotted (IB) for acetylated K268 (Ac-K268), Ac-K293, and total acetylation (ac-K). C. PPARγ acetylation assay of eWAT; DIO mice were treated with vehicle, TPMD, or Rosi (30 mg/kg.bw) for 1 week. Total protein lysates extracted from eWAT were immunoprecipitated (IP) by acetylated lysine (ac-K) antibody conjugated agarose beads and IB for PPARγ. HSP90 was used as a loading control of total protein input. D. In vitro PPARγ acetylation assay in which acetylated PPARγ was co-immunoprecipitated from HEK293 cells transfected with PPARγ and CBP. Co-immunoprecipitated PPARγ was incubated with Sirt1 in the presence of TPMD or Rosi at 37C for 2 hr. Acetylation status was detected by immunoblotting using antibody against Acetyl-lysine. E-F. Luciferase assays of 3xPPRE (E) and Adipsin promoter (F) reporters for deacetylation-resistant PPARγ P495L in HEK293 cells. G–H. PPARγ acetylation in HEK293 cells co-transfected with CBP, SirT1, and either wild-type (WT) or mutant PPARγ as indicated, in the presence or absence of TPMD (30 μM). Total protein lysates were immunoprecipitated with ac-K antibody–conjugated agarose beads and IB for PPARγ. GAPDH was used as a loading control of total protein input. G: PPARγ mutants C313W, R316W, and E371R. H: PPARγ mutants F292C/G370C and F275C/G374C. I. Luciferase assay of 3xPPRE reporter in PPARγ-treated HEK293 cells treated with TPMD or Rosi in the presence or absence of PPARγ antagonist GW9662 (GW). Data were presented as the ratio of treatment in the presence of GW over treatment in the absence of GW. Data are the mean± SEM. * P < 0.05, ** P <0.01, and *** P <0.001.

Next, our observation that TPMD-2 directly interacts with R316 and E371 (R288 and E343 in PPARγ1, respectively) prompted us to hypothesize that R316 and E371 may be involved in TPMD’s induction of PPARγ deacetylation. To test this hypothesis, we generated two mutant forms of PPARγ2 R316W and E371R as well as a third mutant on an adjacent amino acid (C313R), which is predicted to impede the binding of TPMD-2. Treatment of TPMD failed to suppress the acetylation of these PPARγ mutants (Fig. 7G), suggesting that TPMD binding to these amino acids is critical for its acetylation inhibitory activity.

K268 and K293 (K240 and K265 in PPARγ1) are located in the H2-S1 and Ω loop, respectively. The Ω loop is uniquely modulated by TPMD binding (Figs. 6E and 6H). For instance, the Cα of K293 moves 11.3 Å toward the alternative ligand binding pocket of PPARγ LBD (Fig. 6H). We hypothesized that the conformations of β-sheets and Ω loop rendered by TPMD binding may impact their accessibility to acetylation/deacetylation enzymes for PPARγ acetylation regulation. We therefore generated a double mutant form of PPARγ F275C/G374C, in which both F275 and G374 were converted to cysteines that are predisposed to form a stable disulfide bond to stabilize the β-sheets (Fig. S8C) (Montanari et al., 2020), and examined its acetylation status. The PPARγ F275C/G374C mutant was markedly less acetylated compared to WT PPARγ (Fig. 7H). Similarly, a second pair of double mutant form of PPARγ F292C/S370C that is also expected to form a stable disulfide bond to stabilize the Ω loop showed less acetylation than WT PPARγ (Fig. 7H). When cells expressing either double mutant PPARγ (F275C/G374C or F292C/S370C) are treated with TPMD, we observed the further diminishment of PPARγ acetylation (Fig. 7H). These results indicate that formation of disulfide bridge and hence stabilization of the β-sheets and Ω loop keep PPARγ at the state of deacetylation, which is further facilitated by TPMD binding.

Lastly, given the alternative binding pattern of TPMD to PPARγ LBD revealed in our crystal structure, we employed a potent PPARγ antagonist GW9662, which binds and forms an irreversible covalent bond with the canonical PPARγ LBD (IC_50_ 3.3 nM) (Leesnitzer et al., 2002), to block the access to the canonical binding site. As expected, GW9662 significantly diminished the activation of 3xPPRE reporter by Rosi; in contrast, TPMD retained the reporter activity (Fig. 7I), reinforcing the unique mode of TPMD binding to PPARγ.

## Discussion

In this study, we have identified a derivative of a novel chemical scaffold, TPMD, that binds to PPARγ in a unique mode and acts as a weak PPARγ agonist. We further discovered TPMD as a first-in-class small molecule that induces the deacetylation of PPARγ. TPMD improves insulin resistance and glucose tolerance in mouse models of obesity. Importantly, unlike TZDs, which cause severe undesirable effects such as weight gain, adiposity and fluid retention (Choi et al., 2014; Nesto et al., 2004), TPMD reduces body weight and adiposity instead, and is shorn of TZD-associated water retention, cardiac hypertrophy, and bone remodeling. Together, these results demonstrate that TPMD improves insulin sensitization and safety.

Recent studies have shown that PPARγ PTMs are responsible for insulin sensitivity while the undesirable side effects associated with TZD use are likely mediated through the classical full agonism actions of PPARγ (Choi *et al*., 2010; Kroker and Bruning, 2015b; Lazar, 2018). It was recently reported that genetic mimicking of PPARγ deacetylation (2KR) dissociates TZD’s metabolic benefits from its adverse effects (Kraakman *et al*., 2018); however, it is unknown whether PPARγ deacetylation on its own can promote metabolic benefits while sparing any undesirable side effects and if so whether it is achieved through the selective activation of PPARγ target genes. Our studies on TPMD have provided insights to the above unknowns. First, we discovered that TPMD promotes PPARγ deacetylation both *in vitro* and *in vivo* and that PPARγ deacetylation is indispensable for TPMD-induced PPARγ activity. Second, consistent with the phenotypic improvements on insulin sensitivity and other metabolic function at tissue and organismal levels, TPMD treatment preferentially increases the expression of genes associated with insulin sensitivity (Choi *et al*., 2010) in the adipose tissues. Moreover, TPMD treatment augments the expression of thermogenic genes corresponding with the preservation of BAT function, the brown remodeling of WAT, and the enhanced expenditure of organismal energy. On the other hand, unlike Rosi, TPMD shows little or no induction on the expression of genes involved in heart failure or hypertrophy, including β*-Mhc*, *Nppb*, *Bnp*, and *Acta1*, in the heart, on osteogenic/bone forming genes (*Osteocalcin*, *Col1a1*, *Alp*, *Spp1*) in the bone or bone marrow, or on adipogenic genes (*Pparg2*, *Cfd*, *aP2*, *Perilipin*, *Adipoq*) in the adipose tissue, which are associated with PPARγ agonism. Our study therefore indicates that pharmacological deacetylation of PPARγ by TPMD selectively activates the PPARγ target genes to decouple the beneficial metabolic improvements from classical agonism-associated adverse effects.

The co-crystal structure revealed a unique binding mode of TPMD to PPARγ, with three TPMD molecules bound to a single PPARγ LBD: one in the canonical binding site, one in the alternate binding site, and one on the AF-2 surface. First, while the interaction within the canonical binding site is known to stabilize the AF-2 surface for co-activator recruitment to mediate classical PPARγ full agonistic activation by TZDs (Kroker and Bruning, 2015b), which also leads to activation of genes involved in adiposity as well as in many of undesirable side effects (Choi *et al*., 2010), TPMD binding alters the conformation of the canonical site as by Rosi binding, likely resulting in a weak agonism and hence lessening/elimination of side effects. Second, a second TPMD molecule binds to the alternative site binding by extending deeply into branch III in the alternative binding site to engage and stabilize β2 and β3 strands and the Ω loop, which is strikingly different from other PPARγ ligands as they extend to the opening of branch II in the alternative binding site (Jang *et al*., 2019b; Jang *et al*., 2019c). Interaction with the PPARγ alternative binding site is believed to be associated with insulin sensitization, lack the pro-adipogenic effect of TZD, and mediate no classical PPAR signaling (Arifi et al., 2023; Hughes *et al*., 2014). Third, one TMPD molecule binds to the coactivator-binding site of the AF-2 surface, a site normally occupied by the SRC-1 LXXLL motif when co-crystallized with the PPARγ LBD in the absence of TPMD. We wish to point out that it is currently unclear whether the finding of the third TPMD binding in a crystal structure reflects any in vivo functional consequence; however, it raises the possibility that TPMD may impact PPARγ activity by binding to the AF-2 surface and thereby impeding coactivator recruitment. Together, all the interactions with PPARγ LBD by three TPMD molecules hence likely combine to contribute to the metabolic benefits by TPMD while avoiding adverse side effects.

Importantly, the binding of the second TPMD with the alternative binding site of PPARγ LBD provides insight into the mechanism by which TPMD binding results in PPARγ deacetylation. The amino acids K268 and K293 that are subject to acetylation are within the H2-S1 loop and Ω loop that form the outer portion of the alternative binding site of PPARγ LBD. The second TPMD primarily makes multiple interactions with several amino acids in H3 (S317 and R316, as S289 and R288 in PPARγ1) and the β turn between β2 and β3 strands (E371, as E343 in PPARγ1). Although these amino acids are positioned within the more inner part of the alternative binding pocket, these amino acids appear critical for TPMD-induced deacetylation as our site-directed mutagenesis studies indicate that the PPARγ mutants of R316W and E371R are refractory to TPMD for deacetylation. Moreover, the interactions of TPMD-2 with H3 and with the β turn between the β2 and β3 strands make the inner part of the alternative binding pocket to become more compact and stabilized likely through an induced fit and conformational selection mechanism (Shang and Kojetin, 2021; Siclari and Gardner, 2021). Similarly, the interactions of Ω loop with TPMD-2 move Ω loop inward, rendering the Ω loop more stable yet still with significant flexibility. The Ω loop and β-sheets have been proposed to serve as anchors for enzymes that modify PPARγ. For incidence, this region was reported as dock sites for kinases (e.g., CDK5) for S273 phosphorylation (Montanari *et al*., 2020; Mottin et al., 2015). We envision that the flexible stabilization of the β-sheets and Ω loop rendered by TPMD alters the docking of acetylases including CBP and/or deacetylases including SIRT1 for the acetylation status of PPARγ. In supporting this notion, PPARγ mutants predicted to stabilize the Ω loop (F292C/S370C) and β-sheets (F275C/G374C) via the formation of disulfide bonds exhibit a decrease in the acetylation of PPARγ. The decline in the acetylation of PPARγ by TPMD could be due to diminished occupancy of PPARγ acetylases or heightened recruitment of the deacetylase SIRT1 to PPARγ. Our observation that TPMD increases the deacetylation of pre-acetylated PPARγ in the presence of SIRT1 *ex vitro* (Fig. 7D) suggests that TPMD promotes the recruitment of SIRT1 for PPARγ deacetylation.

Although both PPARγ dephosphorylation at S273 (Banks et al., 2015; Choi *et al*., 2010; Choi *et al*., 2011) and deacetylation at K268/K293 (Kraakman *et al*., 2018) were reported to promote insulin sensitization, key differences exist between PPARγ dephosphorylation and PPARγ deacetylation. First, mice with 2KR KI are resistant to weight gain, fluid retention, heart hypertrophy, and bone loss when treated with the TZD drug Rosi (Kraakman *et al*., 2018), whereas mice carrying the dephosphorylation mimetic (S273A) are still susceptible to full adverse effects of Rosi (Hall et al., 2020). Second, 2KR mice, but not S273A KI mice, are protected against diet-induced obesity (Hall *et al*., 2020; Kraakman *et al*., 2018) as 2KR promotes white-to-brown adipose conversion and increases thermogenesis (Kraakman *et al*., 2018). Significantly, TPMD, as the first-in-class small molecule inhibitor of PPARγ acetylation, retains the gist of metabolic benefits in insulin sensitizing and browning as observed in 2KR mice treated with TZD drug but circumvents the undesirable side effects of TZD drugs. Consequently, targeting PPARγ deacetylation could provide a favorable therapeutic option to counter against diabetes, obesity, and their comorbidities.

### Limitations of Study

While we discovered the first-in-class small molecule TPMD that induces PPARγ deacetylation, as a prototype, TPMD is a ligand of PPARγ of rather weak binding affinity with an IC_50_ at ∼5μM. A potent PPARγ ligand of IC_50_ at sub-μM or even nM with the ability for deacetylation induction will be desirable for *in vivo* pharmacokinetics and pharmacodynamics studies as well as for in-depth efficacy studies. Structure-activity-relationship and structure-metabolism-relationship studies will need to be employed to guide the synthesis and optimization of more potent TPMD derivatives that are capable of potently binding PPARγ LBD and inducing PPARγ deacetylation induction but also possess optimal *in vivo* bioavailability for oral administration of the drug, a necessary step for PPARγ deacetylation as a potential anti-diabetic therapy.

## Figure legends

**Figure S1: TPMD protects pancreatic beta cells against ER stress.**

A. Cell viability of INS-1 cells that were treated with TPMD at concentrations as indicated in the presence of Tm (0.1 μg/mL) for 72 h, determined by intracellular ATP assay. B. Live-cell phase-contrast images in INS-1 cells treated as in A. C. Immunoblotting of apoptotic markers in INS-1 cells that were treated with or without Tm (0.3 μg/ml) in the presence of TPMD at indicated concentrations or DMSO for indicated time points. Cleaved caspase-3 and PARP were determined by Western blotting. β actin was used as the loading control. D. Immunoblotting of ER stress markers in INS-1 cells that were treated with or without Tm (0.3 μg/ml) in the presence of TPMD at indicated concentrations or DMSO. β actin was used as a loading control. E-J. qRT-PCR analysis of relative mRNA expression levels of ER stress genes in INS-1 cells that were treated with or without Tm (0.3 μg/ml) in the presence of TPMD at indicated concentrations or DMSO. Data are presented as mean ± SEM. * P < 0.05, ** P <0.01, and *** P <0.001. ns for no significance.

**Figure S2: TPMD shows weak activity in adipocyte differentiation.**

A. 3T3-L1 adipocyte differentiation under the indicated treatment of TPMD or Rosi for last 4 days; Oil Red O staining for lipid accumulation at indicated concentrations. B. Primary mouse adipocyte differentiation with SVF isolated from eWAT under the indicated treatment of TPMD or Rosi for last 4 days. C. qRT-PCR analysis of relative mRNA expression levels of adipogenic genes in cells differentiated from SVF isolated from eWAT under the indicated treatment of TPMD or Rosi for last 4 days as in Fig. S2B. Data are presented as mean ± SEM.* P < 0.05, ** P <0.01, and *** P <0.001. ns for no significance.

**Figure S3: TPMD improves insulin sensitivity and energy expenditure.**

A-B. qRT-PCR analysis of mRNA levels of genes related with insulin sensitivity in eWAT (A) and in iWAT (B) from DIO mice treated with TPMD or vehicle. C-D. qRT-PCR analysis of mRNA levels of adipocyte housekeeping genes in eWAT (C) and in iWAT (D) from DIO mice treated with TPMD or vehicle. E-F. qRT-PCR analysis of mRNA levels of lipogenic (E) and gluconeogenic (F) markers in liver tissues from DIO mice treated with TPMD or vehicle. G-H. Oxygen (G) and carbon dioxide (H) consumption levels, normalized to body weight, in diet-induced obese (DIO) mice treated with TPMD or vehicle for 4 weeks. I. Energy expenditure levels, normalized by body weight. J. Respiratory exchange ratio (RER) measurement. K. Activity, expressed as distance traveled (m/mouse). Data are presented as mean ± SEM.* P < 0.05, ** P <0.01, and *** P <0.001.

**Figure S4: TPMD promotes adipose tissue remodeling.**

A-B. Adipocyte size distribution of eWAT (A) and iWAT (B) from DIO mice treated with TPMD or vehicle. C. Immunofluorescent staining of F4/80 in eWAT sections from DIO mice treated with TPMD or vehicle for 4 weeks. D. Quantification of percentage of F4/80 positive cells in eWAT sections as in C. E. qRT-PCR analysis of mRNA levels of genes for M2 anti-inflammatory markers in eWAT from DIO mice treated with TPMD or vehicle. F. qRT-PCR analysis of mRNA levels of genes known for browning/thermogenesis in iWAT from DIO mice treated with TPMD or vehicle. G. qRT-PCR analysis of mRNA levels of browning/thermogenic marker genes in adipocytes (differentiated from wild-type eWAT) under the indicated treatment of TPMD or DMSO. H. Representative images (arrows) of multilocular cells in eWAT sections from DIO mice treated with TPMD. Data are the mean± SEM. * P < 0.05, ** P <0.01, and *** P <0.001.

**Figure S5: TPMD improves insulin sensitivity and comorbidity in Leptin-deficient obese mice.**

A. Timing of i.p. drug administration. B-C. qRT-PCR analysis for mRNA levels of genes known for insulin sensitivity (B) and for adipocyte housing-keeping (C) in iWAT from *ob/ob* mice treated with TPMD or vehicle. D-E. Immunofluorescent staining of F4/80 in iWAT sections from *ob/ob* mice treated with TPMD or vehicle for 4 weeks; representative images (D) of F4/80 staining and quantification of percentage of F4/80 positive cells (E). F. H&E staining of liver sections from ob/ob mice treated with TPMD or vehicle for 4.5 weeks for both lower and higher magnifications. Data are the mean± SEM. * P < 0.05, ** P <0.01, and *** P <0.001.

**Figure S6: TPMD has no obvious metabolic effects in male C57BL/6 mice fed a normal chow diet.**

A. Glucose tolerance test from mice treated with TPMD or vehicle for 3 weeks. Blood glucose levels measured at indicated time points after intraperitoneal injection of glucose (2 g/kg body weight) following 14-h fasting and the AOC. B. Insulin tolerance test performed for mice treated with TPMD or vehicle for 3.5 weeks. Blood glucose levels normalized to basal level at indicated time points after intraperitoneal injection of insulin (0.7 IU/kg body weight) following 6-h fasting and the AOC. C. Body weight of mice treated with TPMD or vehicle for 4 weeks. D. Daily food intake, measured for three consecutive days during the 4^th^ week of treatment. Changes in body weight of ob/ob mice treated with TPMD or vehicle over the period of 4-wk treatment. E. Measurement of PCV in mice treated with TPMD and vehicle. (F-I). Metabolic cage analysis of ND mice treated with TPMD or vehicle after 4-week treatment, Measurement of oxygen (F) and carbon dioxide (G) consumption levels. H. Energy expenditure levels. (I) Respiratory exchange ratio (RER) measurement. (J). Locomotor activity, expressed as distance traveled (m/mouse). Data are the mean± SEM. * P < 0.05, ** P <0.01, and *** P <0.001.

**Figure S7: Additional structural features of the PPARγ LBD in complex with TPMD.**

The 2Fo – Fc electron density map (1.0 σ) of TPMD-1, TPMD-2 (A) and TPMD-3 (B) were displayed at a radius of 2.0 Å to the TPMDs. (C-E) Conformational changes of PPARγ LBD induced by TPMD-1 (C and D) and TPMD-2 (D and E), in comparison with Rosi-bound PPARγ LBD. The overall structure of Rosi-bound LBD is shown in beige ribbons while its protein residues and Rosi are shown in stick representation with carbon atoms depicted in beige. Helix 3 was hidden for better display of TPMDs in Figure B and D. (F-G) Interactions of the Ω loop with TPMD-2 and the β turn between β2 and β3 strands. (H) An overall structure of the PPARγ LBD containing SRC-1. The LBD and SRC-1 are shown in grey and red ribbons respectively. (I) TPMD-3 functions as a lattice contact between two crystallographic neighboring LBD molecules, colored with light blue and light purple respectively. The LBD in light blue is in the same orientation as in Fig. 6A. TPMD-1, 2 and 3 shown in pink sticks belong to the LBD in light blue while TPMD-1^S^, 2^S^ and 3^S^ shown in purple sticks belong to the crystallographically symmetric neighboring LBD in light purple. Rosi-bound PPARγ LBD (beige) in complex with SRC-1 (red) is superimposed with TPMD-bound LBD (blue).

**Figure S8: Effect of TPMD on PPARγ PTMs**

A-A’. Phosphorylation of PPARγ S273 in 3T3-L1 adipocytes under the treatment of TPMD 20 μM or Rosi 10 μM for 6 h and for TNFα 10 ng/ml for 1 h before harvest; pSer273 of PPARγ was detected by Western blotting using PPARγ pSer273-specific antibody (A) and quantified by image-J (A’). B-B’. Immunoblotting (B) and quantification (B’) for phosphorylation status of PPARγ protein at S273 in eWAT from DIO mice treated with TPMD or vehicle for 4. weeks. Data are the mean± SEM and are representative of 3 independent experiments. * p<0.05, ** p<0.01, *** p<0.001, n=4. (**C**) A cartoon representation of PPARγ LBD in complex of TPMD molecules to highlight the Cα of mutated residues in double mutants F275C/G374C (F247C/G346C in PPARγ1) and F292C/S370C (F264C/S342C in PPARγ1). The Cα of F275, G374, F292, S370 are shown in spheres. The Ω loop and H2-S1 loop are highlighted using thicker loops.

## Experimental methods

### Cell Culture and Plasmids

The 3T3-L1 mouse fibroblasts and HEK-293 cells (ATCC, Manassas, VA, USA) were maintained at 37°C in a humidified 5% CO₂ atmosphere. The cells were cultured in Dulbecco’s modified Eagle’s medium (DMEM) containing 4.5 g/L glucose and 10% fetal bovine serum (FBS; HyClone, Logan, UT, USA). 3T3-L1 preadipocytes were differentiated as previously described (Eeda et al., 2019). The isolation and differentiation of primary preadipocytes were performed as previously described(van Harmelen et al., 2005). Lipid accumulation in differentiated 3T3-L1 cells was assessed using Oil Red O staining (Alfa Aesar, Haverhill, MA, USA). For all cell-based assays, the final concentrations of DMSO did not exceed 0.1%. Plasmids were obtained from addgene: pBabe-bleo-human PPARγ2 (#11439), pcDNA-Flag-Pparg2 (#8895), PPRE X3-TK-luc (#1015), pIS1 (#12179), Sirt1 and CBP. Pparg mutants (C313W, R316W, E371R, F292C/G370C, F275C/G374C) were generated by site-directed mutagenesis. HEK-293T cells were transiently transfected with pcDNA-Pparg together with CBP and Sirt1 using Fugene 6 transfection reagent (Promega, Madison, WI, USA).

### In Vitro Competition Binding Assay

Competition binding assays were performed using the PolarScreen™ PPARγ-Competitor Assay Kit (Thermo Scientific, Waltham, MA, USA), following the manufacturer’s instructions. Briefly, compounds at serial 2-fold dilutions were incubated with GST-tagged PPARγ ligand-binding domain (LBD, 34 nM) and Fluormone PPAR Green tracer (9 nM) in a black 384-well assay plate (Corning Glass catalog no. 677). Reactions were carried out at room temperature for 4 h in the dark. Fluorescence polarization (mP) was measured at 485 nm excitation and 535 nm emission using a microplate reader.

### PPAR**γ** Transactivation Reporter Assay

HEK-293 cells were seeded overnight, co-transfected with plasmids expressing human PPARγ2, 3xPPRE-luc and pIS1 (from Addgene) followed by incubation for 16 h. Cells re-seeded in a 384-well plate, incubated for 24 h and then treated with compounds for an additional 24 h. Luciferase activity was analyzed in each well using Dual-Glo or Bright-Glo Luciferase kits from Promega (Madison, WI, USA).

### Animal Studies

C57BL/6J mice and obese ob/ob mice were obtained from Jackson Laboratory (Bar Harbor, ME, USA). All procedures involving animals were approved by the Institutional Animal Care and Use Committee of the University of Oklahoma Health Science Center and Columbia University Medical Center. The mice were housed at 22°C under a 12-hour light/dark cycle with ad libitum access to food and water. All experiments were conducted with age-matched mice. Male C57BL/6J mice were fed a high-fat diet (HFD) (60% kcal from fat, Bio-Serv, NJ, USA) starting at 5-6 weeks of age. After 8 weeks of HFD feeding, the mice were randomly divided into two groups and administered either vehicle (10% DMSO) or TPMD (30 mg/kg body weight in 10% DMSO) once daily via intraperitoneal injection. Obese ob/ob mice, at 8 weeks of age, were also randomly grouped for intraperitoneal injection with either vehicle or TPMD. An intraperitoneal glucose tolerance test (ipGTT) was performed after a 16-hour overnight fast, with blood glucose levels measured at 0, 15, 30, 60, and 120 minutes using a glucometer (OneTouch Ultra 2 Meter) following intraperitoneal administration of glucose at a dose of 1.5 g/kg body weight. An intraperitoneal insulin tolerance test (ipITT) was performed after a 6-hour fast. Mice were injected with human insulin (1 IU/kg body weight, i.p.), and blood glucose was measured at the same time points. Packed cell volume (PCV) was measured with blood drawn from tail on a LW Scientific E8 microhematocrit centrifuge (LW Scientific, Lawrenceville, Georgia, USA) towards the end of the treatment. Body composition, including lean and fat mass, as well as total body weight, was measured using EchoMRI (EchoMRI LLC, Houston, TX, USA).

### RNA extraction and qRT-PCR

Total RNA was isolated from tissues or cells using TRIzol reagents (Life Technologies) and quantified. Two micrograms of RNA were reverse transcribed with oligo(dT) primers (New England Biosystems) using the SuperScript IV kit (Applied Biosystems). qPCR was performed on a CFX96 Touch Real-Time PCR detection system (Applied Biosystems) using SYBR Green mix (Applied Biosystems). The amplification program was as follows: initial denaturation at 95°C for 15 min, followed by 40 cycles of 95°C for 15 s, 60°C for 1 min, and 40°C for 30 s. Relative expression was calculated by the ΔΔCt method normalized to *TBP* mRNA. The sequences of primers used in this study are listed in Supplemental Table 1.

### Protein extraction and immunoblotting analysis

We extracted protein with RIPA buffer containing protease and phosphatase inhibitors (Thermo Fisher Scientific). Lysates were centrifuged at 10,000LJg for 10 min and total protein concentrations in the cell lysate were determined by BCA assay. Equal protein (∼30 µg) was denatured in 5% β-mercaptoethanol, separated on a 4-12% Bis-Tris NuPAGE gels (Invitrogen), and transferred to polyvinylidene-fluoride membrane at 40 V for 1 h at 4°C. Membranes were blocked with 3% BSA in TBS (10 mm Tris-HCl, pH 7.4, 150 mM NaCl) for 1 h at room temperature and probed with primary antibodies followed by HRP-conjugated secondary antibodies (1:5000; Santa Cruz Biotechnology). The primary antibodies were: Pparg (#2430, Cell Signaling), phosphor-Ser-273 PPARγ (bs-4888R, Bioss Antibodies), GAPDH (#5174, Cell Signaling) and UCP1 (14670S, Cell Signaling).

### Immunoprecipitation and Pparg acetylation assays

For cell-based determination of Pparg acetylation, HEK-293 cells were seeded overnight and then transfected with various Pparg mutants, as well as CBP and Sirt plasmids using Fugene 6 (Promega, Madison, WI). 24 hours after transfection, the cells were treated with either TPMD or Rosi for 6 hours. Proteins were then extracted using an acetylation-specific IP buffer (50 mM Tris, pH 7.5, 150 mM NaCl, 1% IGEPAL, 0.05% SDS, 0.25% sodium deoxycholate). Acetylated proteins were purified by incubating the lysates with acetyl-lysine affinity beads (AAC04, Cytoskeleton, Inc.) overnight at 4°C on a rotator. Beads were subsequently washed for four times and boiled in non-reducing sample buffer. For ex vitro acetylation assay, HEK293 cells were seeded overnight and then transfected with Flag-PPARγ and CBP plasmids at a ratio of 2:1 using Fugene 6 (Promega, Madison, WI) for 24 hours. Cells were then harvested using Flag IP buffer (50 mM Tris-HCl [pH 7.9], 150 mM NaCl, 10% glycerol) supplemented with protease inhibitors. Flag-PPARγ protein was purified by incubating the lysates with Anti-Flag Magnetic Agarose Beads (A36797, Thermo Scientific, Waltham, MA, USA) overnight at 4°C on a rotator. Beads were subsequently washed for four times and boiled in non-reducing sample buffer. Deacetylation reactions were performed at 37°C for 2 hours by mixing recombinant SIRT1 protein (#10011190, Cayman Chemical, Ann Arbor, MI) and the IP-purified PPARγ with the indicated compounds (vehicle, TPMD, or Rosi) in 1× reaction buffer containing 50 mM Tris-HCl (pH 8.8), 4 mM MgCl₂, 1 mM dithiothreitol (DTT), 100 µg/mL bovine serum albumin (BSA), and 0.05 mM NAD⁺.

### Histology and immunohistochemistry

Adipose tissue and liver samples were dissected, immediately fixed in 4% paraformaldehyde (Sigma–Aldrich), paraffin-embedded, and sectioned for H&E staining. For immunohistochemistry, sections were stained with UCP1 antibody (ab10983, Abcam). For immunofluorescence staining, paraffin-embedded epididymal (eWAT) and inguinal (iWAT) adipose tissue sections were deparaffinized and incubated overnight at 4 °C with F4/80 antibody (#70076, Cell Signaling). Slides were washed in PBS containing 0.2% Triton X-100 and incubated with Alexa Fluor 488-conjugated secondary antibodies (Jackson ImmunoResearch, PA, USA) and DAPI (0.5 μg/mL) for 1.5 hours at room temperature. Images were acquired using an Olympus Fluoview 1000 laser-scanning confocal microscope (Center Valley, PA, USA) and quantified using ImageJ histogram software.

### Biochemical analysis

Serum insulin (ALPCO, NH, USA), serum cholesterol and triglycerides (Cayman Chem., MI, USA) and free fatty acids (Bioassay system, CA, USA) were determined using ELISA kites according to the manufactures’ instruction.

### Indirect calorimetry measurements

Mice were individually housed in chambers of a Promethion Core Monitoring system (Sable Systems, Las Vegas, NV, USA). After a 1-day acclimation, their oxygen consumption (VO₂), carbon dioxide production (VCO₂), and locomotor activity were recorded continuously for 3 consecutive days. The respiratory exchange ratio (RER) and energy expenditure were calculated using standard equations.

### Cloning, protein expression and purification

Amino acids 195-477 of human PPARγ-1 was subcloned into the T7 *E. coli* expression vector pMCSG7. This construct of the recombinant protein encodes a 29-residue N-terminal tail (MHHHHHHHHHH SSGVDLGTENLYFQ↓SNAM) containing a 10His tag and a TEV cleavage site in front of the starting residue Ala195. The His-tagged plasmid was transformed into BL21(DE3) cells, which were grown at 25 °C, induced with 1 mM isopropyl-β-D-thiogalactoside (IPTG) at A_600_ = 1.0 for 3 hours and then collected. The cells were lysed by emulsifier in buffer A (20 mM Tris-HCl at pH 8.5, 150 mM NaCl, 10% glycerol and 0.1 mM TCEP) containing 5 mM imidazole and 1 mM PMSF. The crude lysate was centrifuged at 40,000 g for 1 h. The supernatant was applied to a HisTrap HP column (Cytiva), which was previously equilibrated with buffer A containing 5 mM imidazole. After the recombinant protein was eluted with gradient increase to buffer A containing 500 mM imidazole, the eluted protein was desalted in buffer A with PD10 desalting column (Cytiva) to remove imidazole. The N-terminal His tag was cleaved with TEV protease at 4 °C overnight. The N-terminal fusion tag and uncleaved material were removed with a HisTrap HP column (Cytiva). The flow-through was applied to a Superdex 200 Increase 10/300 column (Cytiva), which was previously equilibrated with buffer A. Fractions containing the human hPPARγ LBD were pooled and concentrated to 15 mg/ml for crystallization.

### Crystallization, X-ray data collection, structure determination and refinement

Before crystallization, the purified hPPARγ LBD and a LXXLL motif-containing peptide derived from human steroid receptor coactivator-1 (SRC-1) (residues 686-700, RHKILHRLLQEGSPS) were mixed in a ratio of 1:2, with a 5-fold molar excess of the TPMD compound. After overnight incubation, the mixture was crystallized using the hanging drop vapor diffusion method at 20 °C. The crystals for hPPARγ LBD in complex with TPMD grew in drops consisting of 1 μL of protein solution and 1 μL of reservoir solution to equilibrate against 600 μL of reservoir solution 2.14 M sodium malonate pH 7.0 and 3% (v/v) DMSO.

For crystallization of the hPPARγ LBD in complex with SRC-1, a similar protocol was used except that no TPMD was added, and the reservoir contained 67% Tacsimate (pH 7.0) (Jang et al., 2019a).

Diffraction data were collected on 19-ID NYX beamline at Brookhaven National Laboratory. Raw X-ray diffraction data were indexed and integrated using X-ray Detector Software (XDS). Multiple datasets were scaled and merged using XSCALE, followed with anisotropic scaling and truncation by the Staraniso server. Crystal structure was solved by molecular replacement using apo-hPPARγ LBD (PDB: 1PRG) and refined with phenix.refine. Diffraction data statistics are summarized in Table S2.

### Statistical Analysis

All data are presented as the mean ± SEM. Statistical significance was determined using unpaired two-tailed Student’s t-test or ANOVA with multiple comparisons, as appropriate. A p-value < 0.05 was considered statistically significant. For energy expenditure experiments, group effects were analyzed by ANOVA, with adjustments for body mass and interaction effects using a generalized linear model (GLM).

**Table S1.**
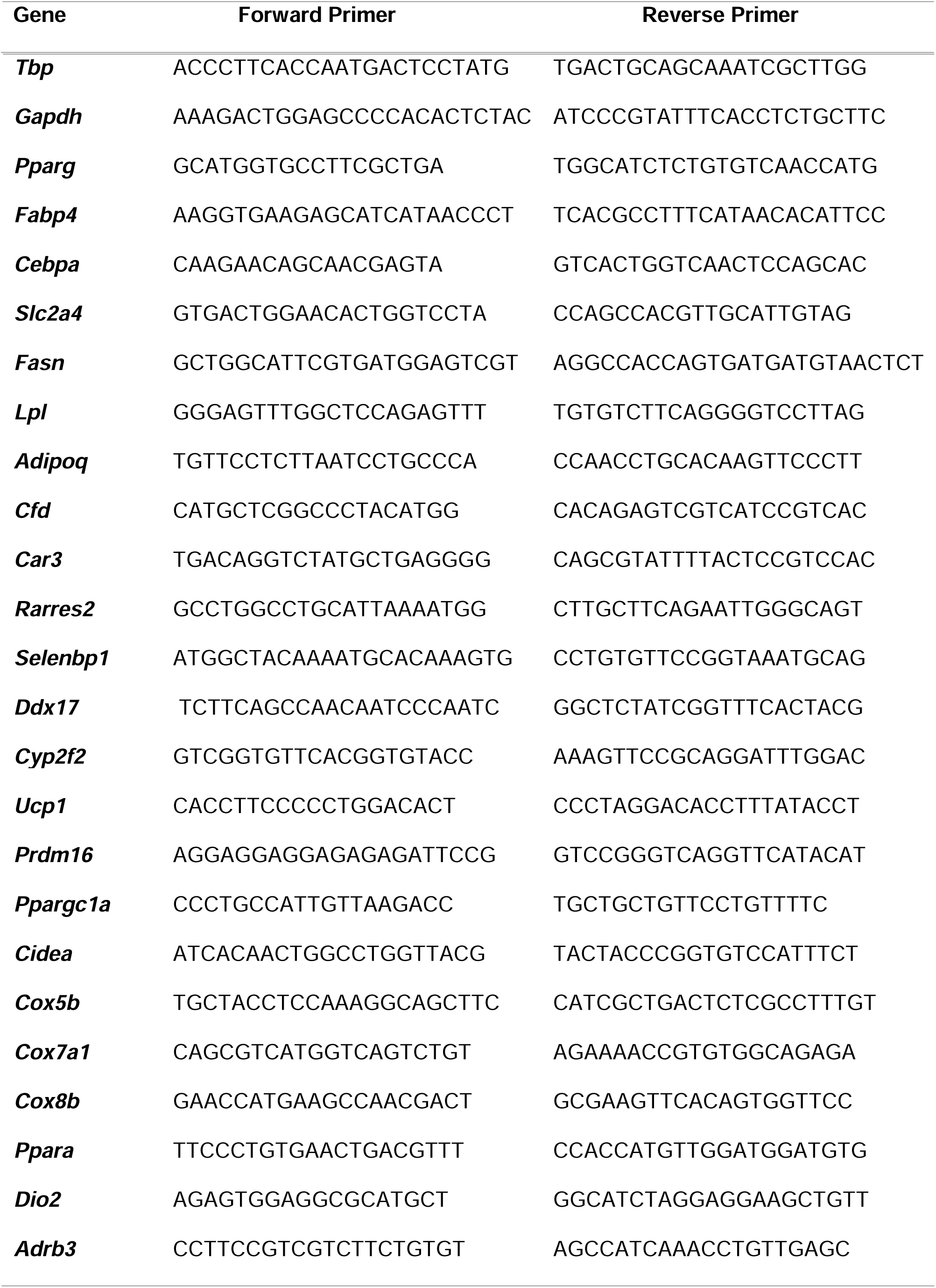
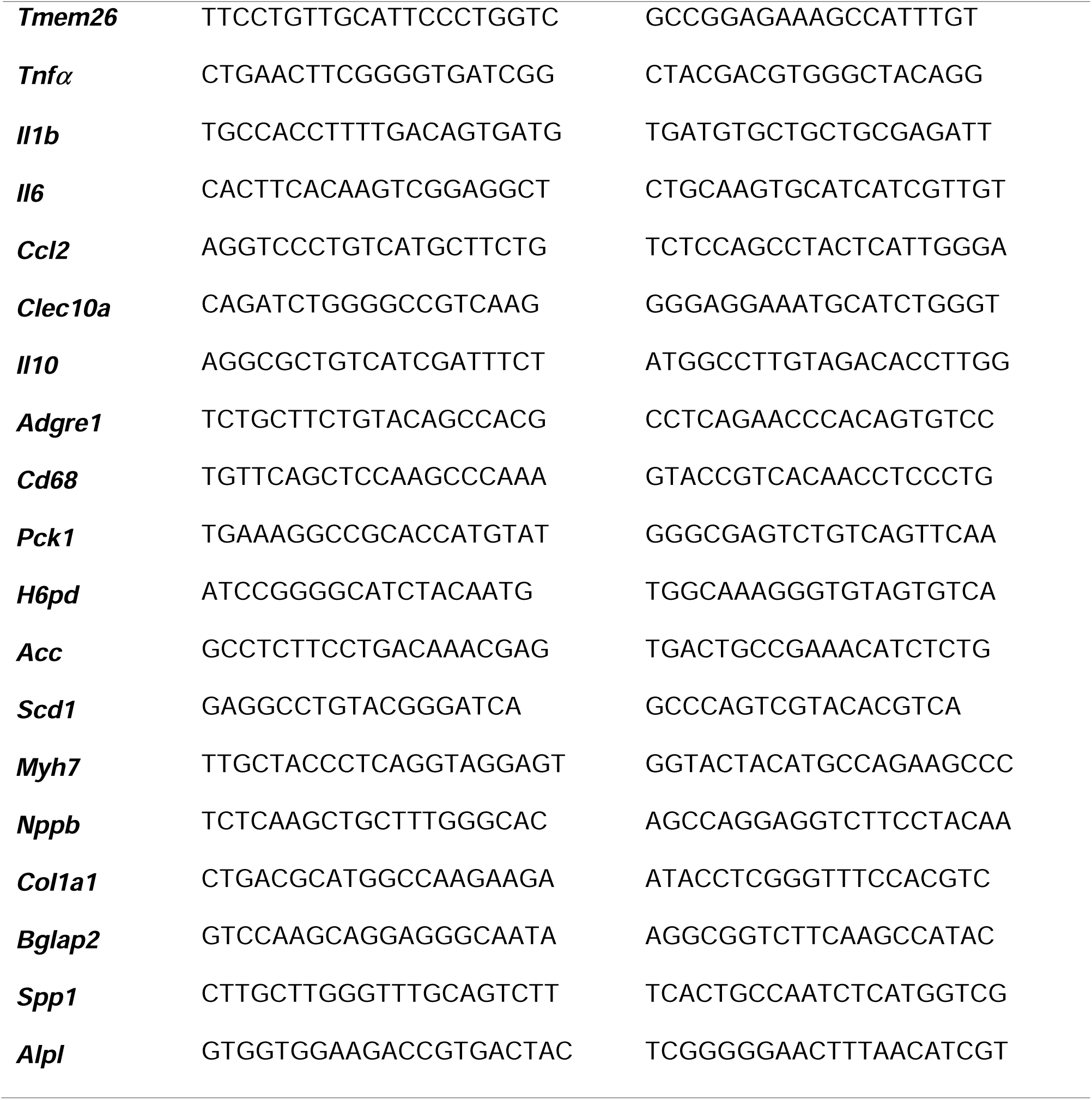
Primers used in this work.

**Table S2.**
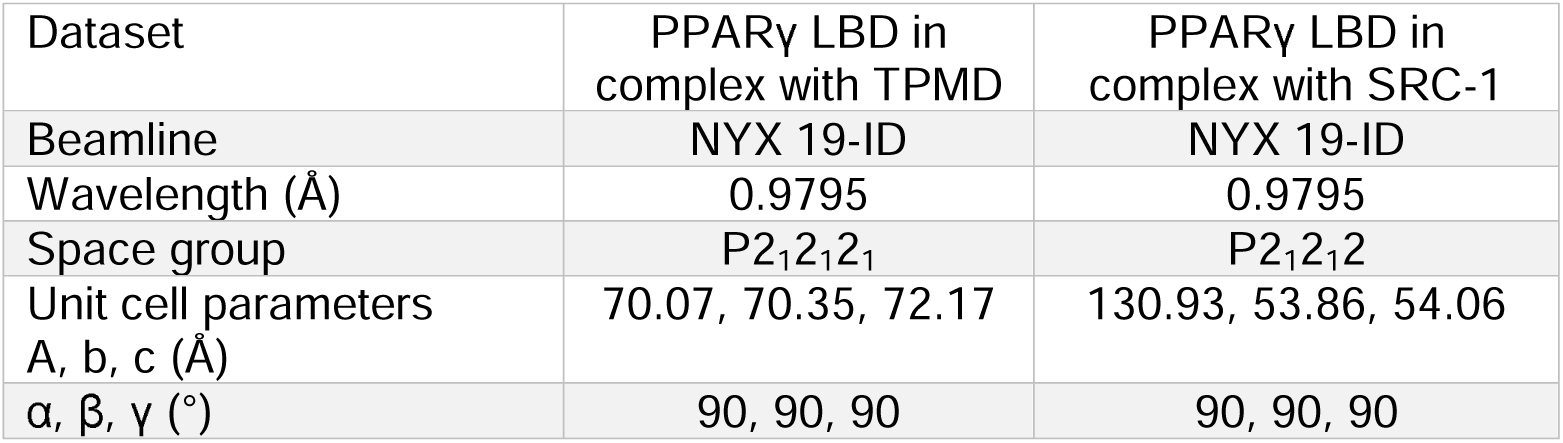
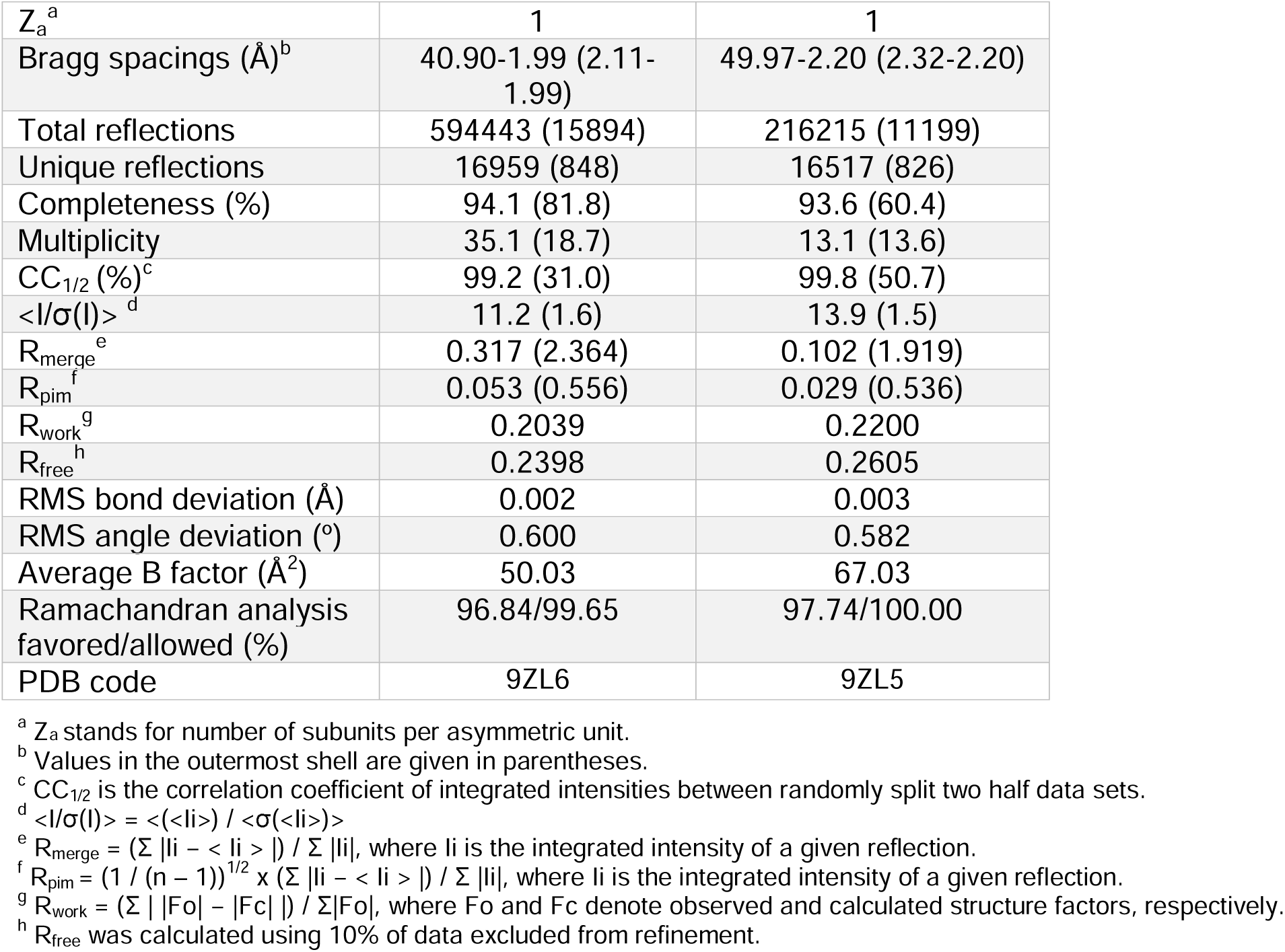
Diffraction Data Statistics.

## Acknowledgments

We thank the Diabetes CoBRE histology and Imaging (5P30GM122744) and Molecular Biology and Cytometry Research (P30CA225520) Cores at OUHSC for assistance on histology and microscopy use. We acknowledge the staff of the NYX beamline at NSLS-II, especially Dr. Kevin P. Battaile, for their support during X-ray diffraction data collection. This work was supported by grants from the NIH R01DK128848 (W.W. and L.Q.), R01DK130174 (W.W.), and R01DK112943 (L.Q.).

## Author Contribution

W.W. and L.Q. conceived and supervised the study, designed the experiments, and obtained funding. D.W. Y.H., S.K., Q.W., O.H-P, and W.W. designed and performed in vitro and in vivo experiments. V.E designed and synthesized chemicals. Z.G designed and performed crystallization experiments. W.A.H. supervised crystallization experiments. All authors processed and interpreted the data. D.W., Z.G., and Y.H. created the figures. W.W., Z.G., and L.Q. wrote the manuscript with editing from D.W, Y.H., H-Y.L., and W.A.H.

## References

Aaron, N., Zahr, T., He, Y., Yu, L., Mayfield, B., Pajvani, U.B., and Qiang, L. (2022). Acetylation of PPARγ in macrophages promotes visceral fat degeneration in obesity. Life Metab 1, 258–269. 10.1093/lifemeta/loac032.

Arifi, S., Marschner, J.A., Pollinger, J., Isigkeit, L., Heitel, P., Kaiser, A., Obeser, L., Höfner, G., Proschak, E., Knapp, S., et al. (2023). Targeting the Alternative Vitamin E Metabolite Binding Site Enables Noncanonical PPARγ Modulation. Journal of the American Chemical Society 145, 14802–14810. 10.1021/jacs.3c03417.

Banks, A.S., McAllister, F.E., Camporez, J.P.G., Zushin, P.-J.H., Jurczak, M.J., Laznik-Bogoslavski, D., Shulman, G.I., Gygi, S.P., and Spiegelman, B.M. (2015). An ERK/Cdk5 axis controls the diabetogenic actions of PPARγ. Nature 517, 391–395. 10.1038/nature13887.

Choi, J.H., Banks, A.S., Estall, J.L., Kajimura, S., Bostrom, P., Laznik, D., Ruas, J.L., Chalmers, M.J., Kamenecka, T.M., Bluher, M., et al. (2010). Anti-diabetic drugs inhibit obesity-linked phosphorylation of PPARgamma by Cdk5. Nature 466, 451–456. 10.1038/nature09291.

Choi, J.H., Banks, A.S., Kamenecka, T.M., Busby, S.A., Chalmers, M.J., Kumar, N., Kuruvilla, D.S., Shin, Y., He, Y., Bruning, J.B., et al. (2011). Antidiabetic actions of a non-agonist PPARgamma ligand blocking Cdk5-mediated phosphorylation. Nature 477, 477–481. 10.1038/nature10383.

Choi, S.S., Kim, E.S., Koh, M., Lee, S.J., Lim, D., Yang, Y.R., Jang, H.J., Seo, K.A., Min, S.H., Lee, I.H., et al. (2014). A novel non-agonist peroxisome proliferator-activated receptor gamma (PPARgamma) ligand UHC1 blocks PPARgamma phosphorylation by cyclin-dependent kinase 5 (CDK5) and improves insulin sensitivity. The Journal of biological chemistry 289, 26618–26629. 10.1074/jbc.M114.566794.

Duan, H., Arora, D., Li, Y., Setiadi, H., Xu, D., Lim, H.Y., and Wang, W. (2016a). Identification of 1,2,3-triazole derivatives that protect pancreatic beta cells against endoplasmic reticulum stress-mediated dysfunction and death through the inhibition of C/EBP-homologous protein expression. Bioorganic & medicinal chemistry 24, 2621–2630. 10.1016/j.bmc.2016.03.057.

Duan, H., Lee, J.W., Moon, S.W., Arora, D., Li, Y., Lim, H.Y., and Wang, W. (2016b). Discovery, Synthesis, and Evaluation of 2,4-Diaminoquinazolines as a Novel Class of Pancreatic beta-Cell-Protective Agents against Endoplasmic Reticulum (ER) Stress. J Med Chem 59, 7783–7800. 10.1021/acs.jmedchem.6b00041.

Duan, H., Li, Y., Arora, D., Xu, D., Lim, H.Y., and Wang, W. (2017). Discovery of a Benzamide Derivative That Protects Pancreatic beta-Cells against Endoplasmic Reticulum Stress. J Med Chem 60, 6191–6204. 10.1021/acs.jmedchem.7b00435.

Eeda, V., Wu, D., Lim, H.Y., and Wang, W. (2019). Design, synthesis, and evaluation of potent novel peroxisome proliferator-activated receptor γ indole partial agonists. Bioorganic & medicinal chemistry letters 29, 126664. 10.1016/j.bmcl.2019.126664.

Evans-Molina, C., Robbins, R.D., Kono, T., Tersey, S.A., Vestermark, G.L., Nunemaker, C.S., Garmey, J.C., Deering, T.G., Keller, S.R., Maier, B., and Mirmira, R.G. (2009). Peroxisome proliferator-activated receptor gamma activation restores islet function in diabetic mice through reduction of endoplasmic reticulum stress and maintenance of euchromatin structure. Molecular and cellular biology 29, 2053–2067. 10.1128/mcb.01179-08.

Ghali, O., Al Rassy, N., Hardouin, P., and Chauveau, C. (2016). Increased Bone Marrow Adiposity in a Context of Energy Deficit: The Tip of the Iceberg? Front Endocrinol (Lausanne) 7, 125. 10.3389/fendo.2016.00125.

Gupta, D., Kono, T., and Evans-Molina, C. (2010). The role of peroxisome proliferator-activated receptor gamma in pancreatic beta cell function and survival: therapeutic implications for the treatment of type 2 diabetes mellitus. Diabetes, obesity & metabolism 12, 1036–1047. 10.1111/j.1463-1326.2010.01299.x.

Hall, J.A., Ramachandran, D., Roh, H.C., DiSpirito, J.R., Belchior, T., Zushin, P.H., Palmer, C., Hong, S., Mina, A.I., Liu, B., et al. (2020). Obesity-Linked PPARγ S273 Phosphorylation Promotes Insulin Resistance through Growth Differentiation Factor 3. Cell metabolism 32, 665–675.e666. 10.1016/j.cmet.2020.08.016.

He, Y., B’Nai Taub, A., Yu, L., Yao, Y., Zhang, R., Zahr, T., Aaron, N., LeSauter, J., Fan, L., Liu, L., et al. (2023a). PPARγ Acetylation Orchestrates Adipose Plasticity and Metabolic Rhythms. Adv Sci (Weinh) 10, e2204190. 10.1002/advs.202204190.

He, Y., Zhang, R., Yu, L., Zahr, T., Li, X., Kim, T.W., and Qiang, L. (2023b). PPARγ Acetylation in Adipocytes Exacerbates BAT Whitening and Worsens Age-Associated Metabolic Dysfunction. Cells 12. 10.3390/cells12101424.

Hotamisligil, G.S., Arner, P., Caro, J.F., Atkinson, R.L., and Spiegelman, B.M. (1995). Increased adipose tissue expression of tumor necrosis factor-alpha in human obesity and insulin resistance. J Clin Invest 95, 2409–2415. 10.1172/JCI117936.

Hughes, T.S., Giri, P.K., de Vera, I.M., Marciano, D.P., Kuruvilla, D.S., Shin, Y., Blayo, A.L., Kamenecka, T.M., Burris, T.P., Griffin, P.R., and Kojetin, D.J. (2014). An alternate binding site for PPARγ ligands. Nature communications 5, 3571. 10.1038/ncomms4571.

Itoh, T., Fairall, L., Amin, K., Inaba, Y., Szanto, A., Balint, B.L., Nagy, L., Yamamoto, K., and Schwabe, J.W. (2008). Structural basis for the activation of PPARgamma by oxidized fatty acids. Nat Struct Mol Biol 15, 924–931. 10.1038/nsmb.1474.

Jang, J.Y., Kim, H., Kim, H.J., Suh, S.W., Park, S.B., and Han, B.W. (2019a). Structural basis for the inhibitory effects of a novel reversible covalent ligand on PPARgamma phosphorylation. Sci Rep 9, 11168. 10.1038/s41598-019-47672-w.

Jang, J.Y., Kim, H., Kim, H.J., Suh, S.W., Park, S.B., and Han, B.W. (2019b). Structural basis for the inhibitory effects of a novel reversible covalent ligand on PPARγ phosphorylation. Scientific reports 9, 11168. 10.1038/s41598-019-47672-w.

Jang, J.Y., Kim, H.J., and Han, B.W. (2019c). Structural Basis for the Regulation of PPARγ Activity by Imatinib. Molecules (Basel, Switzerland) 24. 10.3390/molecules24193562.

Jiang, H., Zhou, X.E., Shi, J., Zhou, Z., Zhao, G., Zhang, X., Sun, Y., Suino-Powell, K., Ma, L., Gao, H., et al. (2020). Identification and structural insight of an effective PPARγ modulator with improved therapeutic index for anti-diabetic drug discovery. Chemical science 11, 2260–2268. 10.1039/c9sc05487a.

Khalaf, K.I., and Taegtmeyer, H. (2012). After avandia: the use of antidiabetic drugs in patients with heart failure. Texas Heart Institute journal 39, 174–178.

Kraakman, M.J., Liu, Q., Postigo-Fernandez, J., Ji, R., Kon, N., Larrea, D., Namwanje, M., Fan, L., Chan, M., Area-Gomez, E., et al. (2018). PPARgamma deacetylation dissociates thiazolidinedione’s metabolic benefits from its adverse effects. The Journal of clinical investigation 128, 2600–2612. 10.1172/JCI98709.

Kroker, A.J., and Bruning, J.B. (2015a). Review of the Structural and Dynamic Mechanisms of PPARgamma Partial Agonism. PPAR Res 2015, 816856. 10.1155/2015/816856.

Kroker, A.J., and Bruning, J.B. (2015b). Review of the Structural and Dynamic Mechanisms of PPARγ Partial Agonism. PPAR research 2015, 816856. 10.1155/2015/816856.

Lazar, M.A. (2018). Reversing the curse on PPARγ. The Journal of clinical investigation 128, 2202–2204. 10.1172/jci121392.

Leesnitzer, L.M., Parks, D.J., Bledsoe, R.K., Cobb, J.E., Collins, J.L., Consler, T.G., Davis, R.G., Hull-Ryde, E.A., Lenhard, J.M., Patel, L., et al. (2002). Functional consequences of cysteine modification in the ligand binding sites of peroxisome proliferator activated receptors by GW9662. Biochemistry 41, 6640–6650. 10.1021/bi0159581.

Lehrke, M., and Lazar, M.A. (2005). The many faces of PPARgamma. Cell 123, 993–999. 10.1016/j.cell.2005.11.026.

Li, D., Zhang, F., Zhang, X., Xue, C., Namwanje, M., Fan, L., Reilly, M.P., Hu, F., and Qiang, L. (2016). Distinct functions of PPARgamma isoforms in regulating adipocyte plasticity. Biochem Biophys Res Commun 481, 132–138. 10.1016/j.bbrc.2016.10.152.

Liu, L., Fan, L., Chan, M., Kraakman, M.J., Yang, J., Fan, Y., Aaron, N., Wan, Q., Carrillo-Sepulveda, M.A., Tall, A.R., et al. (2020). PPARgamma Deacetylation Confers the Anti-Atherogenic Effect and Improves Endothelial Function in Diabetes Treatment. Diabetes. 10.2337/db20-0217.

Montanari, R., Capelli, D., Yamamoto, K., Awaishima, H., Nishikata, K., Barendregt, A., Heck, A.J.R., Loiodice, F., Altieri, F., Paiardini, A., et al. (2020). Insights into PPARγ Phosphorylation and Its Inhibition Mechanism. J Med Chem 63, 4811–4823. 10.1021/acs.jmedchem.0c00048.

Mottin, M., Souza, P.C., and Skaf, M.S. (2015). Molecular Recognition of PPARγ by Kinase Cdk5/p25: Insights from a Combination of Protein-Protein Docking and Adaptive Biasing Force Simulations. J Phys Chem B 119, 8330–8339. 10.1021/acs.jpcb.5b04269.

Nanjan, M.J., Mohammed, M., Prashantha Kumar, B.R., and Chandrasekar, M.J.N. (2018). Thiazolidinediones as antidiabetic agents: A critical review. Bioorganic chemistry 77, 548–567. 10.1016/j.bioorg.2018.02.009.

Nesto, R.W., Bell, D., Bonow, R.O., Fonseca, V., Grundy, S.M., Horton, E.S., Le Winter, M., Porte, D., Semenkovich, C.F., Smith, S., et al. (2004). Thiazolidinedione use, fluid retention, and congestive heart failure: a consensus statement from the American Heart Association and American Diabetes Association. Diabetes Care 27, 256–263. 10.2337/diacare.27.1.256.

Nolte, R.T., Wisely, G.B., Westin, S., Cobb, J.E., Lambert, M.H., Kurokawa, R., Rosenfeld, M.G., Willson, T.M., Glass, C.K., and Milburn, M.V. (1998). Ligand binding and co-activator assembly of the peroxisome proliferator-activated receptor-gamma. Nature 395, 137–143. 10.1038/25931.

Qiang, L., Wang, L., Kon, N., Zhao, W., Lee, S., Zhang, Y., Rosenbaum, M., Zhao, Y., Gu, W., Farmer, S.R., and Accili, D. (2012). Brown remodeling of white adipose tissue by SirT1-dependent deacetylation of Ppargamma. Cell 150, 620–632. 10.1016/j.cell.2012.06.027.

Shang, J., and Kojetin, D.J. (2021). Structural mechanism underlying ligand binding and activation of PPARγ. Structure (London, England : 1993) 29, 940–950.e944. 10.1016/j.str.2021.02.006.

Siclari, J.J., and Gardner, K.H. (2021). Two steps, one ligand: How PPARγ binds small-molecule agonists. Structure (London, England : 1993) 29, 935–936. 10.1016/j.str.2021.08.005.

Spiegelman, B.M. (1998). PPAR-gamma: adipogenic regulator and thiazolidinedione receptor. Diabetes 47, 507–514.

Tontonoz, P., Hu, E., and Spiegelman, B.M. (1995). Regulation of adipocyte gene expression and differentiation by peroxisome proliferator activated receptor gamma. Curr Opin Genet Dev 5, 571–576. 0959-437X(95)80025-5 [pii].

Tontonoz, P., and Spiegelman, B.M. (2008). Fat and beyond: the diverse biology of PPARgamma. Annual review of biochemistry 77, 289–312. 10.1146/annurev.biochem.77.061307.091829.

Tran, K., Li, Y., Duan, H., Arora, D., Lim, H.Y., and Wang, W. (2014). Identification of small molecules that protect pancreatic beta cells against endoplasmic reticulum stress-induced cell death. ACS chemical biology 9, 2796–2806. 10.1021/cb500740d.

van Harmelen, V., Skurk, T., and Hauner, H. (2005). Primary culture and differentiation of human adipocyte precursor cells. In Human cell culture protocols, (Springer), pp. 125–135.

Weisberg, S.P., McCann, D., Desai, M., Rosenbaum, M., Leibel, R.L., and Ferrante, A.W., Jr. (2003). Obesity is associated with macrophage accumulation in adipose tissue. J Clin Invest 112, 1796–1808.

