## Supplementary figures and images for "Small molecule inhibitor of PPARγ acetylation promotes insulin sensitization and browning of white adipose tissue with improved safety"

### Supplemental files

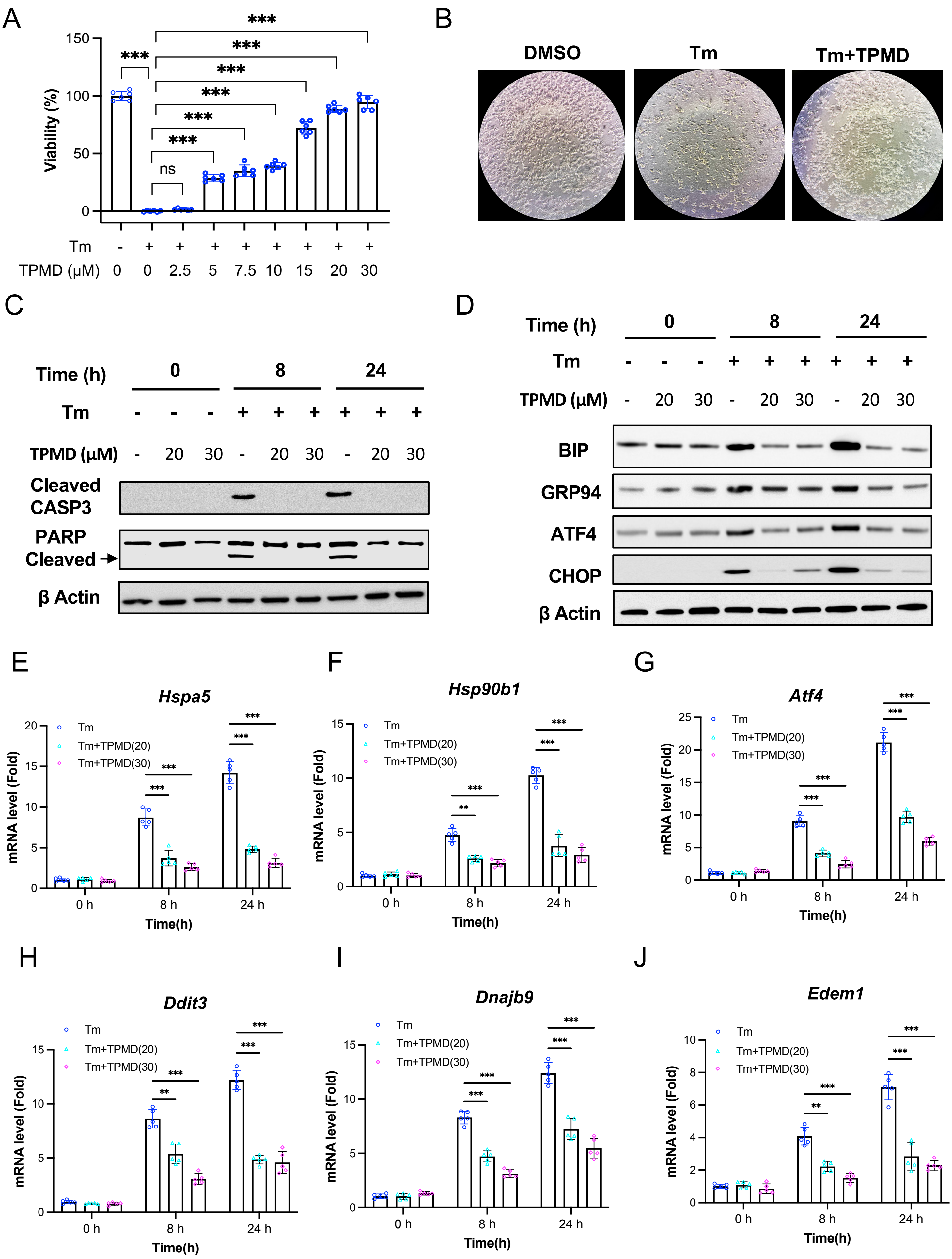

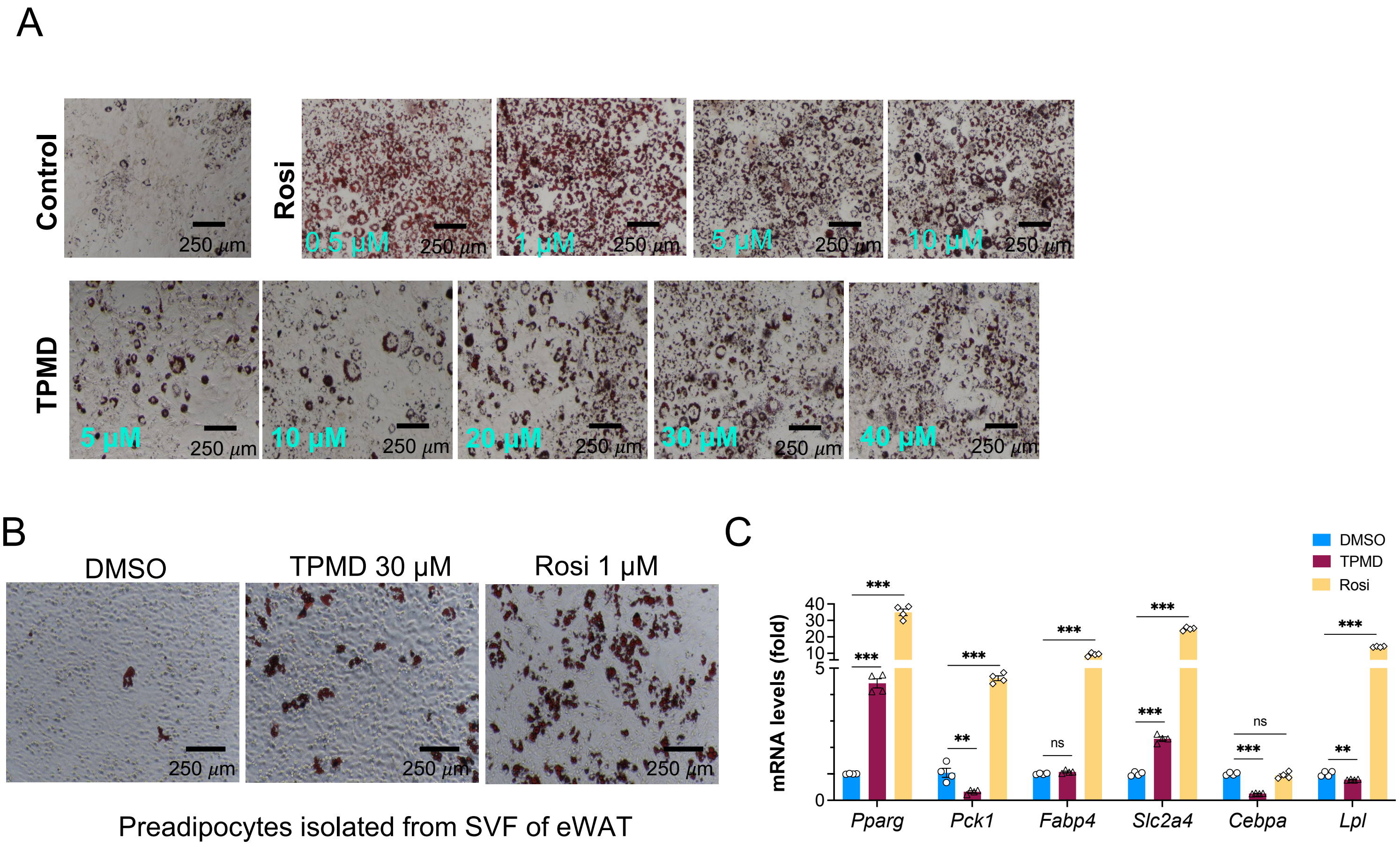

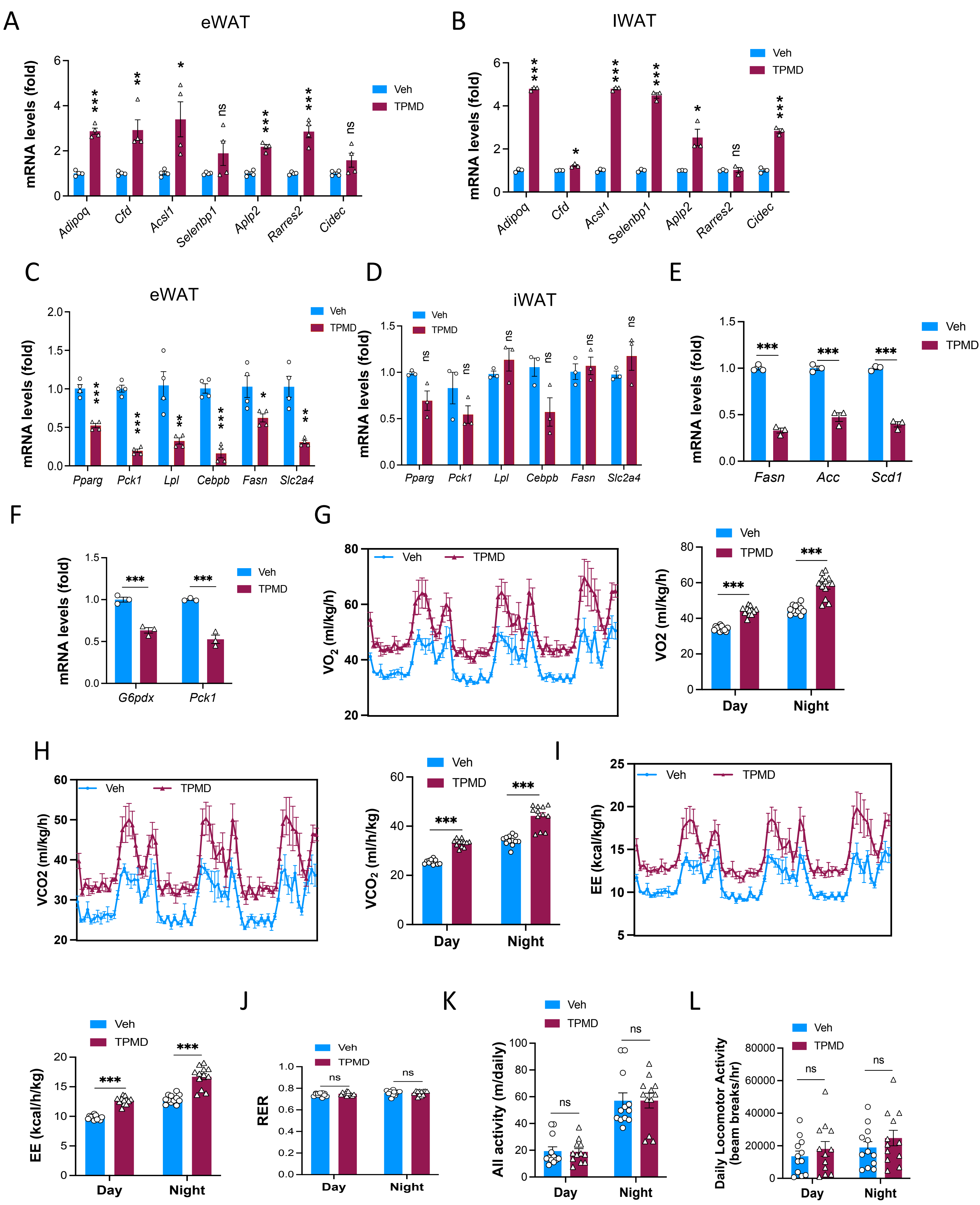

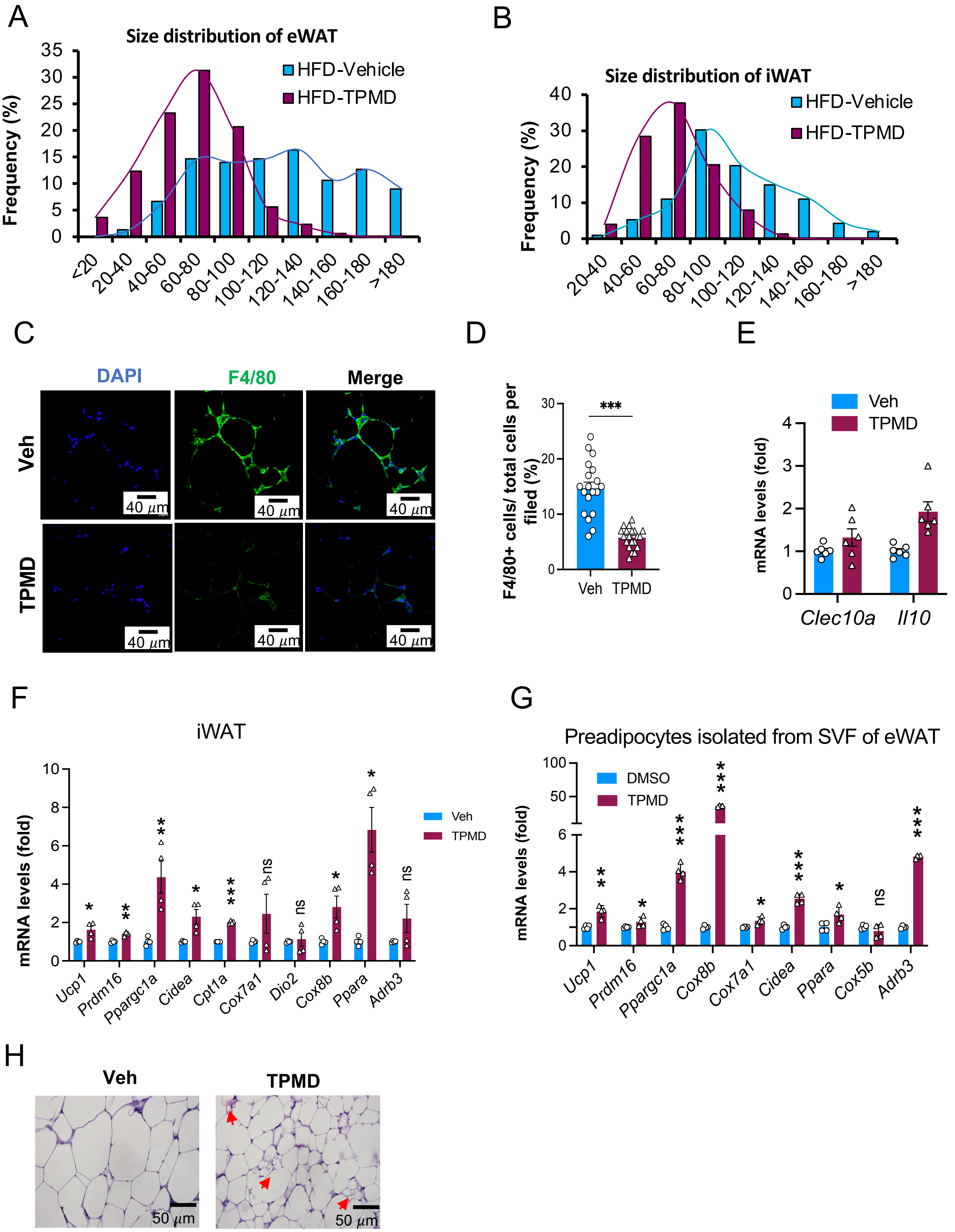

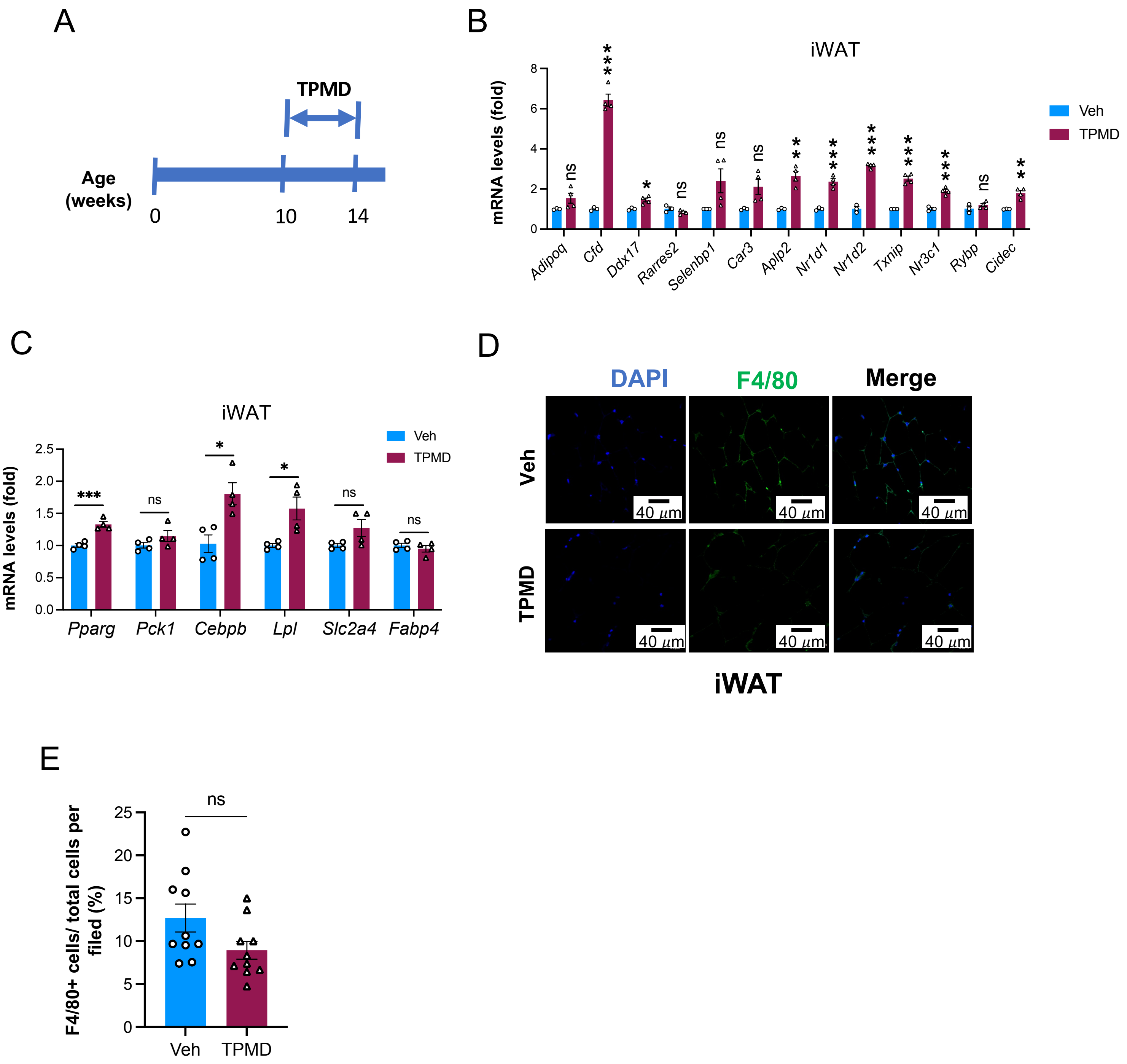

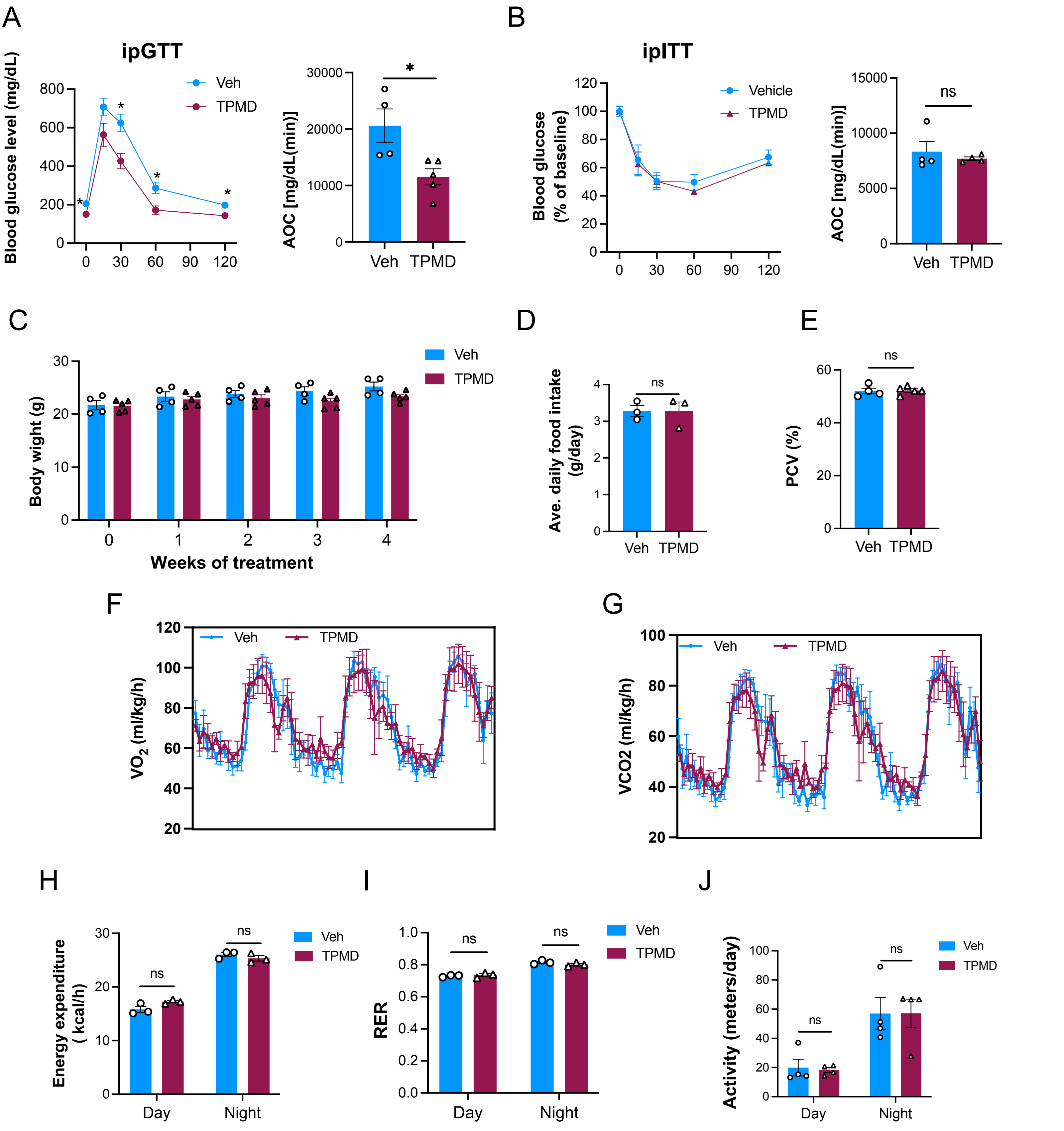

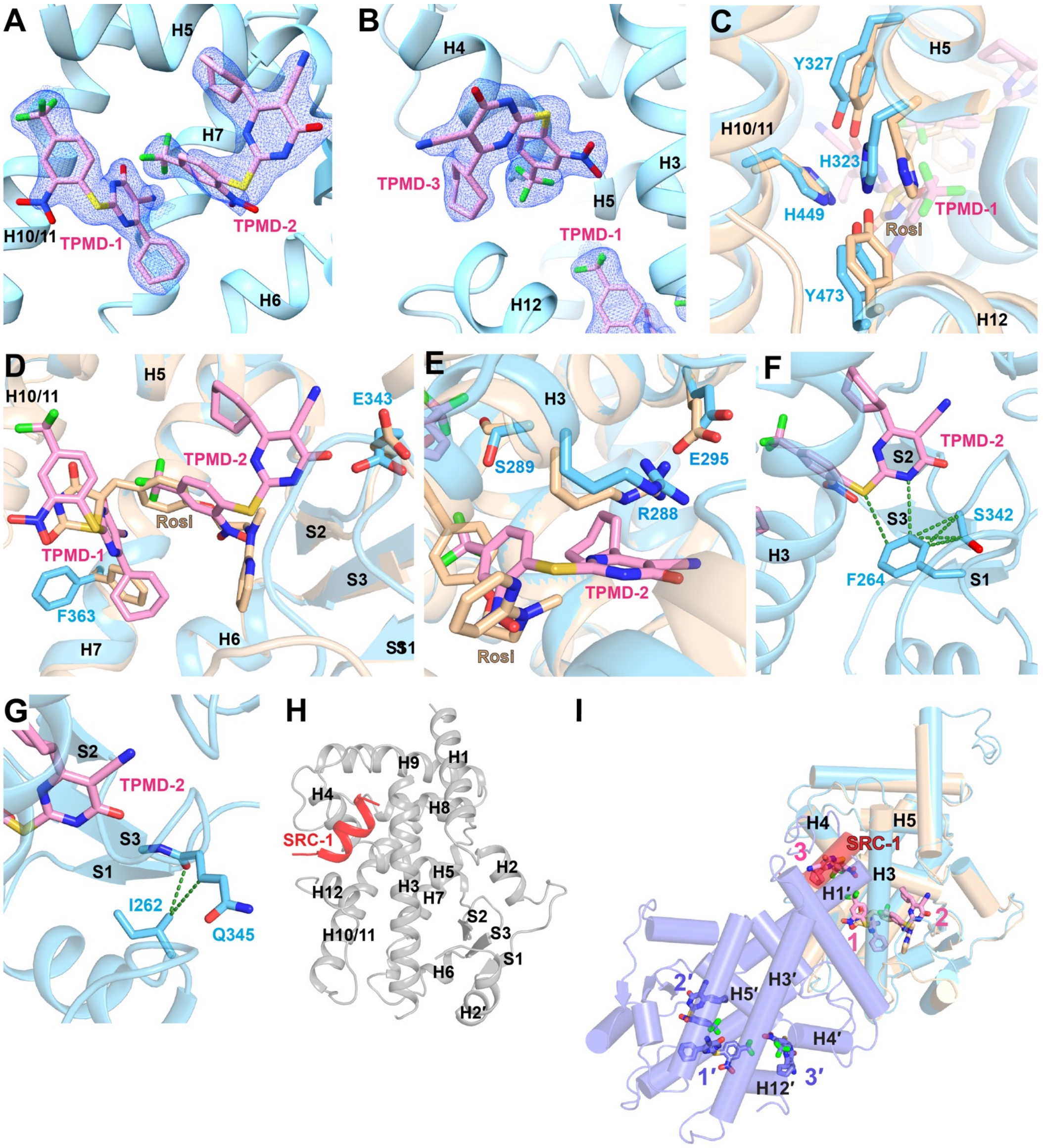

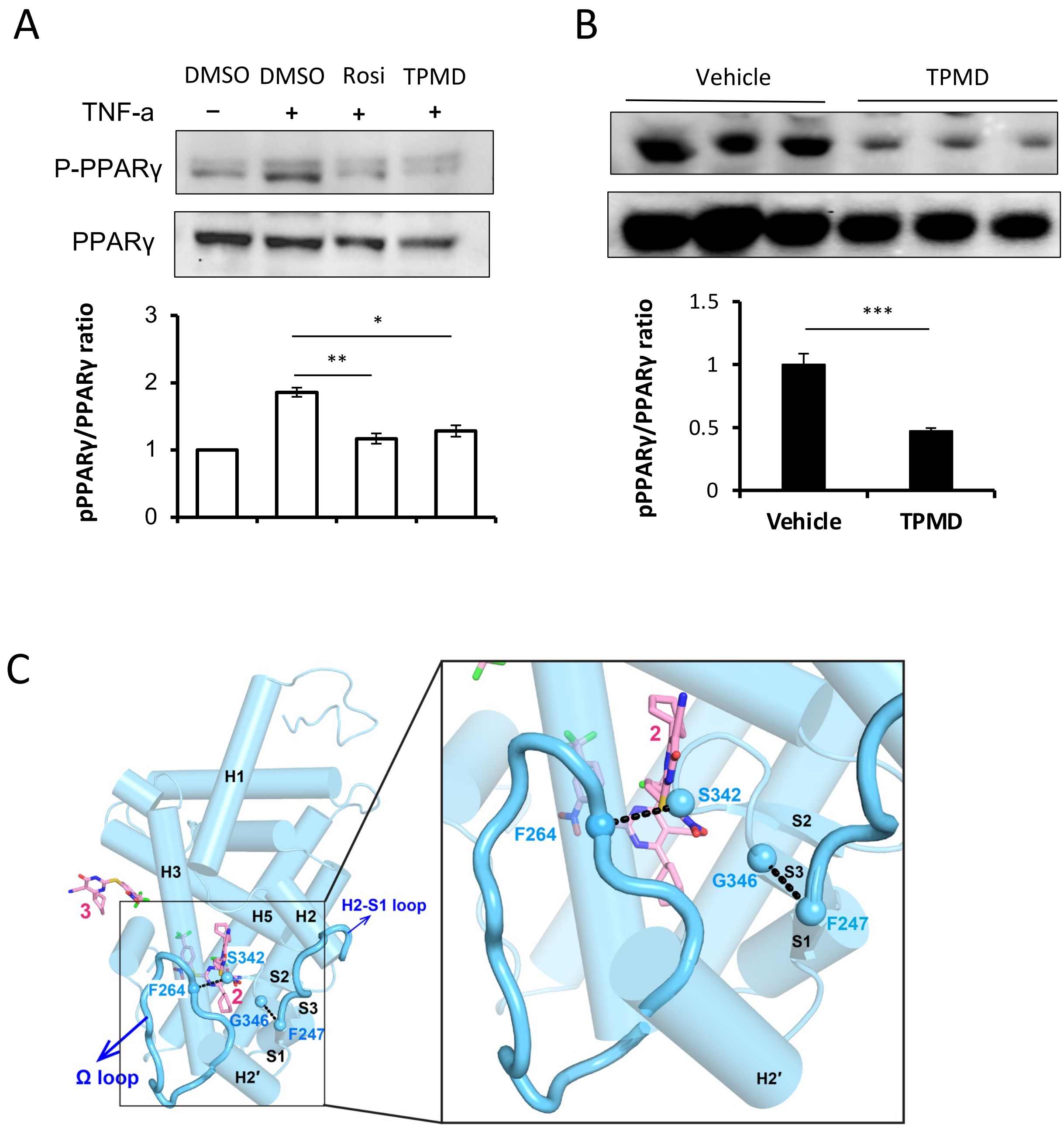
